# Ribosome Molecular Aging Shapes Translation Dynamics

**DOI:** 10.64898/2026.03.08.710403

**Authors:** Jordy F. Botello, Lifei Jiang, Peter J. Metzger, Troy J. Comi, Aya A. Abu-Alfa, Qiwei Yu, Margaret S. Ebert, Minkyu Lee, Lennard W. Wiesner, Maya Butani, Claire J. Weaver, Andrej Košmrlj, Ileana M. Cristea, Clifford P. Brangwynne

## Abstract

Cellular homeostasis relies on continual renewal of cellular components, yet some complexes like ribosomes persist for long periods, raising the question of whether extended molecular age impacts functional fidelity. Here, we introduce a spatiotemporal mapping strategy to resolve biomolecular life stages, and show that intracellular ribosome aging alters translational dynamics at specific transcripts. Molecularly aged ribosomes exhibit impaired elongation at basic amino acid-rich sequences, leading to increased pausing, premature termination, and ribosome collisions. By profiling ribosomal RNA modifications, we find that molecular aging increases the collision propensity of specific ribosome subpopulations. Consistent with our findings, enrichment of aged ribosomes in cells amplifies molecular age-dependent translation defects. *In vivo* labeling of ribosomes in aged *C. elegans* demonstrates that molecularly aged ribosomes shape translational dynamics during organismal aging. These findings identify ribosome molecular age as a determinant of translational dynamics, and link molecular aging of a core gene-expression complex to organismal aging.

**HIGHLIGHTS:**

- A pulse-chase labeling strategy enables mapping subcellular demographics of macromolecular complexes in space and time.
- Molecular aging of ribosomes drives differential mRNA translation and shapes elongation dynamics.
- The collision propensity of specific ribosome subpopulations increases with molecular age.
- Older ribosomes shape translation dynamics during organismal aging.

## INTRODUCTION

Cellular homeostasis depends on a dynamic balance between the synthesis and degradation of biomolecules, enabling the continual renewal of functional components while ensuring the removal of damaged or obsolete ones.^1,2^ Although cellular biomolecules are typically short-lived, with most mRNAs typically decaying on timescales of hours,^3^ and most proteins usually remaining for one to two days in mammalian cells,^4,5^ a distinct subset exhibits remarkable stability. Notable examples include nuclear pore complexes,^6^ chromatin-associated proteins such as histones^7^ and lamins,^8^ and cellular machines like ribosomes,^9,10^ which display substantially longer lifetimes than the proteome at large, with reported values exceeding 100 hours *in vivo*. This contrast is even more pronounced in post-mitotic tissues, where numerous cellular components undergo minimal turnover and can persist for months to years.^11–13^ Despite extensive progress in elucidating pathways governing biomolecular synthesis and degradation, the mechanisms and functional implications of exceptional molecular longevity remain poorly understood.

The persistence of certain long-lived biomolecules is particularly striking, given that cellular turnover is expected to prevent the accumulation of defective components by promoting their timely clearance. These expectations parallel everyday experience with non-living materials, where structural and chemical transformations gradually develop, as with the formation of cracks in rubber, or the slow rusting of iron, which ultimately cause systemic dysfunction in man-made machines. Consistent with this picture, biomolecules within living cells can develop progressive alterations, including oxidation,^14^ crosslinking,^15^ hydrolysis,^16^ or aggregation.^17^ Such changes could gradually alter their folding, binding affinities, or collective mechanical properties, imparting a temporal dimension to their molecular (dys)function. Indeed, the accumulation of these diverse forms of chemical damage in long-lived proteins increases with organismal age,^18^ suggesting their role as a type of ’molecular clock’.^19^

The concept of progressive physicochemical decline of biomolecules is particularly interesting in the context of large biomolecular machines, such as the ribosome. Ribosomes are essential molecular machines conserved across all domains of life, and represent a striking example of a long-lived, essential cellular complex, whose structural and functional integrity is central to maintaining proteostasis in physiology and disease.^20–23^ Assembled within the nucleolus through a complex, high-flux biogenesis pathway that produces thousands of ribosomes every minute in proliferating cells,^24–26^ this process can consume up to 60% of cellular ATP in eukaryotes,^27,28^ an immense metabolic cost that may explain the ribosome’s exceptional stability. However, such prolonged persistence could also render them vulnerable to cumulative molecular damage,^9^ progressively eroding their functional fidelity over time.

Alterations in ribosome biogenesis and turnover, including aging-associated declines in these processes,^29–31^ coincide with reduced functional fidelity.^32–34^ These defects include altered elongation dynamics, such as increased ribosome pausing at regions rich in basic amino acids, which limits the synthesis of basic proteins, particularly those enriched for RNA- and DNA-binding functions, ultimately contributing to mRNA-protein decoupling and proteostasis collapse.^32,34^ In line with these observations, ribosomal component damage is increasingly recognized as a hallmark of aging and age-associated disorders,^35,36^ potentially arising from the extended lifespans of these complexes within cells. Although age-related functional decline is systemic, encompassing epigenetic,^37^ transcriptomic,^38^ and proteomic alterations,^39,40^ these observations suggest that shifts in molecular demographics, i.e. the age distribution of ribosomes or other cellular components, could shape the functional landscape of the aging cell. However, the lack of methods capable of probing native cellular assemblies over extended molecular timescales has hindered efforts to link molecular age with molecular function.

In this manuscript, we address this gap in the study of molecular aging by developing a method to track biomolecular life stages, revealing that molecular age shapes the functional dynamics of the ribosome. Mechanistically, molecular aging disrupts elongation dynamics, preferentially at regions rich in basic amino acids, leading to increased pausing, premature termination, and collision propensity. Molecular aging increases the collision propensity of specific ribosome subpopulations, defined by their ribosomal RNA modification state, whose depletion improves the translation of proteins rich in basic amino acids, suppresses molecular age-dependent collisions, and enhances resistance to ribotoxic stress. These translation defects can be amplified within the steady-state translatome by engineering imbalances in ribosome age demographics. Finally, we perform *in vivo* labeling of aged ribosomes in aging *C. elegans*, providing evidence that ribosome molecular aging shapes translation dynamics during organismal aging. These findings demonstrate that ribosomal aging remodels proteostasis and cellular homeostasis, highlighting specific ribosome subpopulations as potential therapeutic targets for age-related disease.

## RESULTS

### A system for mapping ribosome spatiotemporal compositional dynamics

To characterize the spatiotemporal trajectories of ribosome composition and function over extended timescales in living cells, we developed a modular pulse-chase labeling strategy based on HaloTag, a protein tag that enables irreversible covalent labeling of proteins with defined ligands.^41^ Homozygous endogenous tagging in U2OS cells of surface-accessible ribosomal proteins from the large (RPL10A) and small (RPS17) subunits (Figure S1A-E), preserved their incorporation into actively translating ribosomes (Figure 1A, Figure S1L,M). We then systematically evaluated HaloTag ligands for intrinsic fluorescence, compatibility with high-affinity antibody-based purification, and bio-orthogonality to avoid interference from endogenous molecules, a common limitation of biotin-based immunopurification strategies for protein purification.^42,43^ HaloTag-TMR conjugates supported live-cell imaging and were specifically recognized by antibodies (Figure 1B, Figure S1K), enabling selective purification of various cellular complexes in native conditions, including nuclear pre-ribosomal subunits (Figure S2E), cytoplasmic mature ribosomes (Figure S2A-D,F,G), and spliceosomes (Figure S2H).

**Figure 1.**
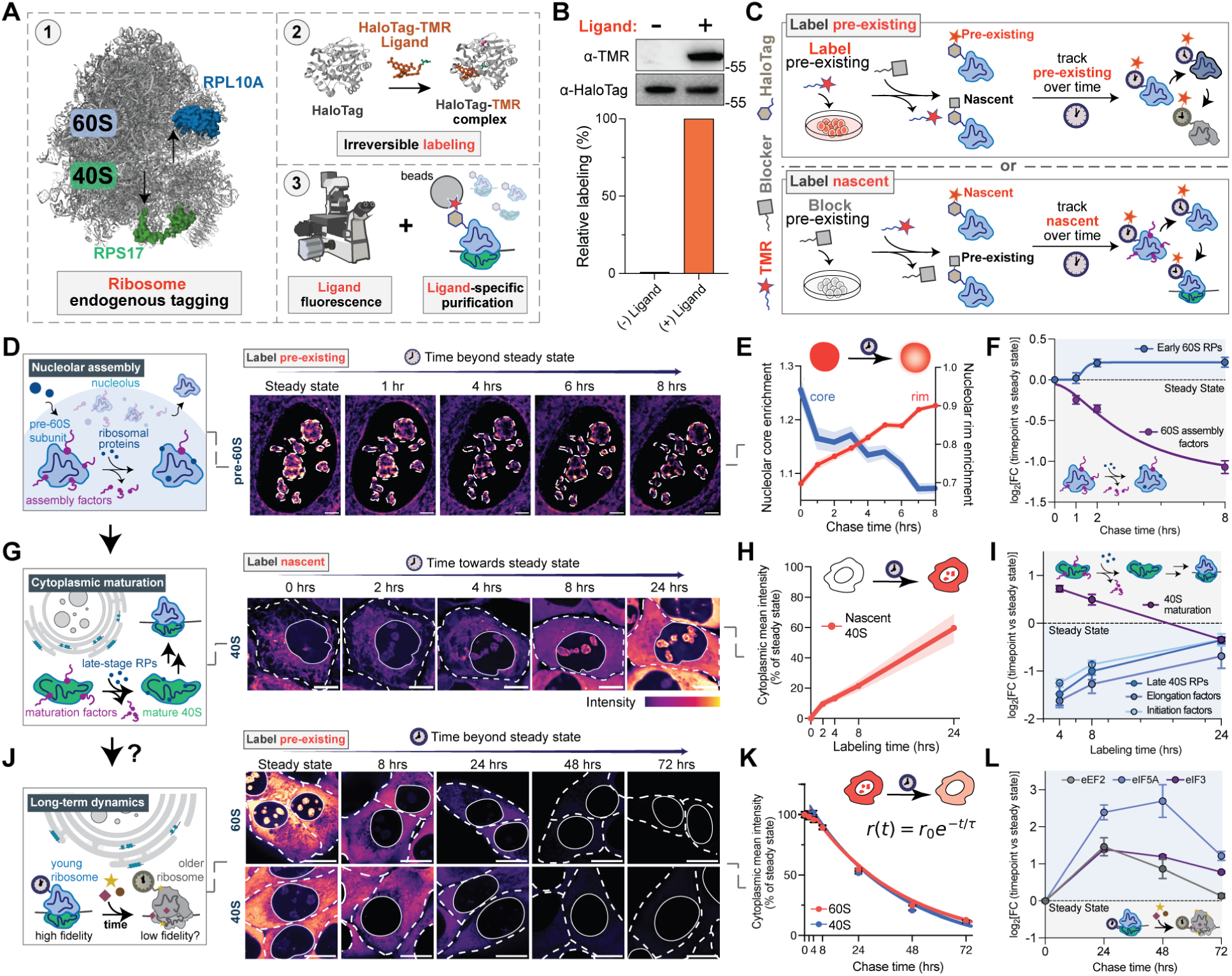
A system for mapping ribosome spatiotemporal compositional dynamics. (A) Strategy for mapping ribosome composition in space and time (PDB: 5AJ0, 6U32). (B) Western blot and quantification of selective anti-TMR detection of TMR-labeled HaloTag-RPL10A relative to total HaloTag-RPL10A. (C) Labeling strategies for tracking pre-existing and nascent molecules over time. (D) Schematic overview of the nucleolar assembly pathway of pre-60S ribosomal subunits, and time-resolved imaging of pre-existing nucleolar pre-60S particles through HaloTag-RPL10A. Scale bar = 5 μm. (E) Quantification of pre-existing pre-60S dynamics in the nucleolus (mean ± S.E.M., n ≥ 700 nucleoli). (F) Dynamics of nuclear pre-60S interactions quantified by immunopurification-mass spectrometry (IP-MS) of time-resolved intermediates (mean ± S.E.M., n = 3). (G) Schematic overview of cytoplasmic maturation of 40S ribosomal subunits and time-resolved imaging of nascent 40S subunits through RPS17-HaloTag. Scale bar = 10 μm. (H) Quantification of nascent cytoplasmic 40S biogenesis relative to steady-state (mean ± S.E.M., n ≥ 379 cells). (I) Dynamics of cytoplasmic nascent 40S interactions quantified by IP-MS of time-resolved intermediates (mean ± S.E.M., n = 3). (J) Schematic overview of long-term cytoplasmic dynamics of mature ribosomal subunits and time-resolved imaging of pre-existing 60S and 40S subunits during dilution and turnover. Scale bar = 15 μm. (K) Quantification and fit of dilution and turnover dynamics of pre-existing 60S and 40S subunits in the cytoplasm relative to steady-state (mean ± S.E.M., n ≥ 57 cells). (L) Dynamics of cytoplasmic pre-existing 40S interactions quantified by IP-MS of time-resolved intermediates (mean ± S.E.M., n = 3).

To resolve distinct stages of the ribosome life cycle across short and extended molecular timescales, we developed complementary HaloTag-TMR pulse-chase labeling schemes to track both pre-existing and nascent ribosome populations (Figure 1C). To track the dynamics of pre-existing populations, the pre-existing HaloTag pool is first saturated with the TMR ligand, followed by ligand washout and addition of a non-fluorescent blocker ligand to prevent further labeling (‘Label-block’, Figure 1C, top). We validated this approach by monitoring the dynamics of nuclear pre-60S ribosomal subunits via HaloTag-RPL10A (Figure 1D,E), whose assembly pathway in human cells is structurally defined.^44^ Time-resolved live-cell imaging revealed the progressive migration of pre-60S intermediates from the nucleolar core toward the periphery, consistent with the outward flux characteristic of nucleolar ribosome assembly.^45,46^ Purification of nuclear intermediates at defined time points, followed by proteomic analysis, recapitulated the expected maturation trajectory, including progressive ribosomal protein incorporation and loss of assembly factor interactions (Figure 1F). Together, these results show that our HaloTag-based strategy accurately tracks pre-existing pre-60S subunits through their expected maturation trajectories.

To track the dynamics of newly synthesized populations, the pre-existing HaloTag pool is first saturated with a blocker, followed by washout and timed HaloTag-TMR labeling (‘Block-label’, Figure 1C, bottom). We applied this approach to track the maturation of nascent ribosomal subunits and their progression into translationally active complexes in the cytoplasm. Time-resolved live-cell imaging at defined intervals enabled visualization and quantification of nascent subunit flux into the cytoplasm (Figure 1G,H). Subsequent purification of cytoplasmic fractions at corresponding time points revealed the expected compositional trajectory of subunit maturation, with early time points enriched for cytoplasmic maturation factors and depleted of translation initiation and elongation factors, progressively shifting toward steady-state interaction profiles consistent with acquisition of translational competence (Figure 1I and Figure S2I). Together, these results demonstrate that this strategy faithfully captures the expected spatiotemporal and compositional dynamics of nascent cytoplasmic ribosomes.

We next asked whether this approach could be extended beyond ribosome maturation to monitor the dynamics of mature ribosomes and their molecular aging over extended timescales. Using the label-block strategy, we tracked pre-existing cytoplasmic ribosomes over hours to days. Time-resolved imaging showed a steady decrease in the concentration of pre-existing ribosomes over time, which was well fitted by an exponential decay function, with a decay timescale of τ = 33 ± 1 hours for labeled 40S, and τ = 30 ± 1 hours for labeled 60S (Figure 1J,K). To understand the origin of these dynamics, we consider that the time rate of change of the ribosome concentration, 𝑟, should be governed by 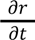 = σ − 𝑘 𝑟 − µ𝑟, where σ is the ribosome production rate, *k*_𝑑𝑒𝑔_ is the degradation rate, and µ is the dilution rate due to cell growth. For the label-block approach (i.e. σ_𝑙𝑎𝑏𝑒𝑙_ = 0 after t = 0), the concentration of labeled ribosomes would be given by 𝑟_𝑙𝑎𝑏𝑒𝑙_ (𝑡) ∼ 𝑒 ^−*t*/τ^, where 1/τ = 𝑘 _𝑑𝑒𝑔_ + µ. By incorporating measurements of the dilution rate µ obtained from quantifying cell growth, we found that 60S and 40S ribosomes exhibit comparable degradation-to-dilution ratios, i.e. *k*_𝑑𝑒𝑔_/µ ∼ 1 (Figure S3A).

To examine whether ribosomes of different ages exhibit distinct functional states, we performed proteomic analysis of cytoplasmic ribosomes isolated at defined molecular ages; we reasoned that if aged ribosomes are functionally distinct, this should manifest in different proteomic compositions. Over multi-day timescales, ribosomes exhibited extensive remodeling of the landscape of interactions with core translation machinery, including translation elongation factors eEF2 and eIF5A, as well as the translation initiation complex eIF3 (Figure 1L). Together, these results demonstrate that our strategy can be used to dissect spatiotemporal stages of the ribosome life cycle, revealing that ribosome interactions with the translational machinery evolve in a molecular age-dependent manner.

### Ribosome molecular aging drives differential mRNA translation

To determine whether the functional landscape of ribosomes is molecular age-dependent, we implemented a molecular age-selective ribosome profiling (Ribo-seq) approach in which footprints from age-specific ribosomes were purified using the label-block strategy and analyzed by deep sequencing (Figure 2A, Figure S3D,E). We performed this analysis for both large (60S) and small (40S) ribosomal subunits, as these dissociate after each round of translation and may thus follow distinct molecular aging trajectories. Consistent with the approximately exponential decay (τ ≅ 30 hrs) of aged ribosomes, ribosomes at least 24 hours old (T_24_) accounted for 54.5 ± 7.6% (40S) and 51.7 ± 6.6% (60S) of the steady-state population, whereas ribosomes at least 72 hours old (T_72_) accounted for 12.7 ± 2.6% (40S) and 10.2 ± 1.3% (60S) (mean ± SD, Figure 2B,C). Polysome profiling of aged (T_72_) versus steady-state ribosomes revealed a pronounced redistribution toward the monosome fraction relative to polysome and free subunit fractions, consistent with age-dependent alterations in translational dynamics (Figure 2D).

**Figure 2.**
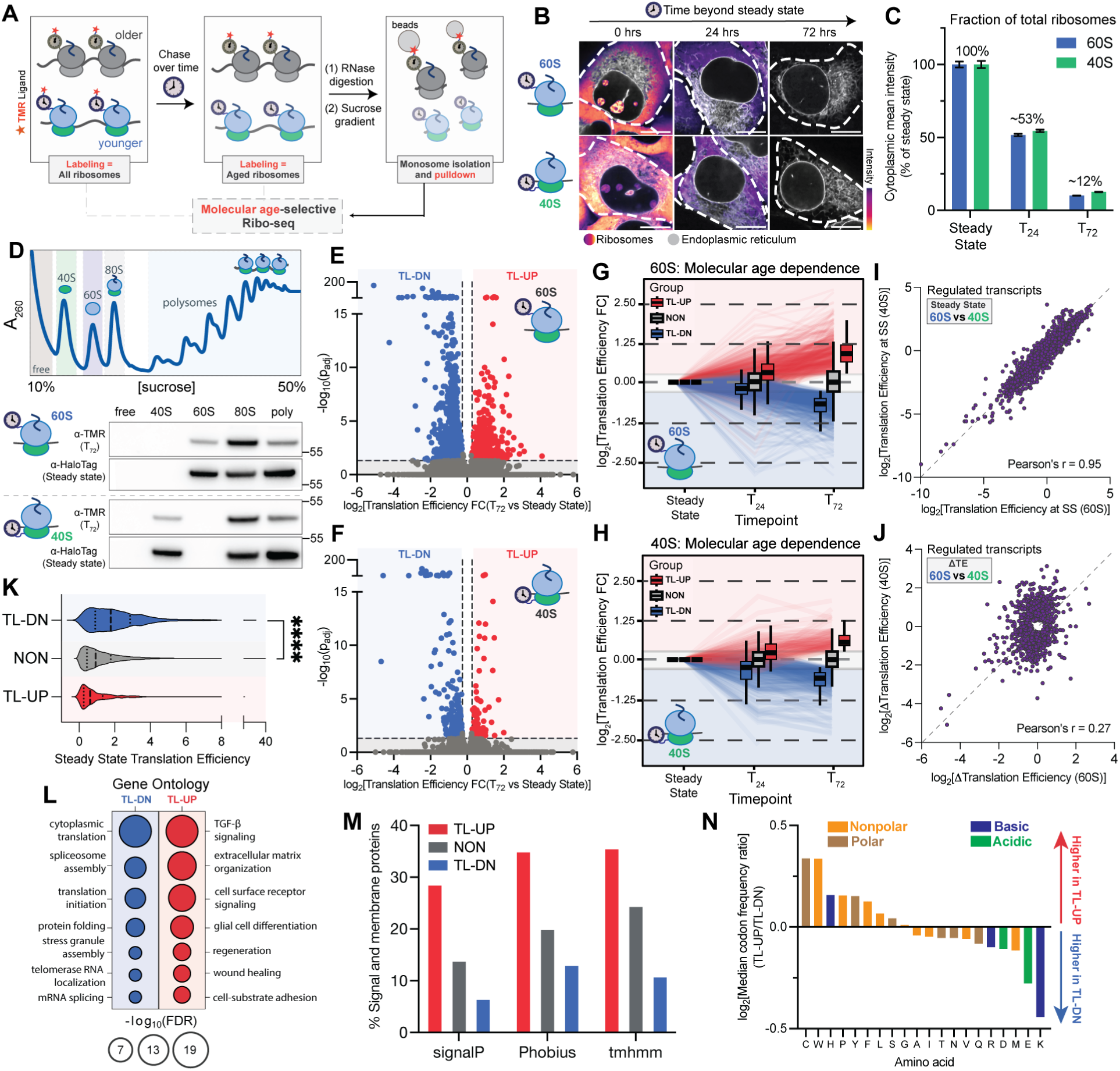
Ribosome molecular age drives differential mRNA translation. (A) Schematic representation of molecular age-selective Ribo-seq experiments. (B) Time-resolved imaging and (C) quantification of ribosome demographics studied, with the endoplasmic reticulum labeled with emiRFP670-SEC61B. Scale bar = 10 μm (mean ± S.E.M, n ≥ 57 cells). (D) Representative polysome profile and western blot analysis of distribution of aged (TMR, T_72_) and steady-state (HaloTag) 60S and 40S ribosome demographics over different fractions. (E,F) Volcano plot of age-dependent translation efficiencies for aged 60S (n = 1,408 regulated transcripts, TL-DN = 881, TL-UP = 527) and 40S (n = 334 regulated transcripts, TL-DN = 236, TL-UP = 98), adjusted p<0.05, FC > 1.2 fold. (G,H) Molecular age-dependent trajectories of TL-UP, NON and TL-DN transcripts for both aging 60S and 40S. (I) Correlation between steady-state translation efficiencies for the union of transcripts regulated by both aged 60S and aged 40S. (J) Correlation between the change in translation efficiencies for the union of transcripts regulated by both aged (T_72_) 60S and aged (T_72_) 40S. (K) Comparison of steady state translation efficiencies between TL-DN and NON regulated transcripts (p < 0.0001, two-tailed Mann-Whitney test) (L) Gene Ontology (Biological Process) terms for the combined set of transcripts regulated by aged 60S and aged 40S subunits (adjusted *P* < 0.05). (M) Percent of signal and membrane proteins encoded by transcripts regulated by aged 60S according to multiple algorithms. (N) Ratio of median codon frequencies between TL-UP and TL-DN transcripts comprising the combined set regulated by aged 60S and aged 40S subunits.

Molecular age-selective Ribo-seq of steady-state, T_24_, and T_72_ ribosomes uncovered widespread changes in translation efficiency (TE) across numerous transcripts, defined as the ratio of ribosome footprint abundance to mRNA abundance, indicating the ribosome occupancy of an mRNA relative to its expression level (Figure 2E,F). These changes occurred progressively over time (Figure 2G,H), consistent with functional remodeling due to molecular aging. Interestingly, rather than a uniform loss of translational activity, we observed distinct classes of transcripts whose translation efficiency was upregulated (TL-UP), downregulated (TL-DN), or non-regulated (NON) for both aged 60S and 40S ribosomes. Although the translation efficiencies for regulated transcripts at steady-state are highly correlated between the 60S and 40S datasets (Pearson’s *r* = 0.95, Figure 2I), the molecular age-dependent changes in translation efficiencies for these transcripts are only weakly correlated (Pearson’s *r* = 0.28, Figure 2J), suggesting that the two subunits follow distinct molecular aging trajectories. Consistent with the reduced polysome association of aged ribosomal subunits (Figure 2D), aged ribosomes preferentially lose translation on transcripts that are highly translated at steady state (Figure 2K). Turnover rates across ribosome age demographics did not differ significantly, (Figure S3B), and cell cycle distributions were comparable across conditions (Figure S3C), ruling out these variables as contributors to the observed translation differences.

To determine whether these transcript classes share common biological features, we next examined their functions. Gene Ontology (GO) analysis of the combined regulated transcript set revealed clear functional divergence between transcript classes (Figure 2L). Transcripts translated more efficiently by aged ribosomes are enriched for processes linked to cell surface receptor signaling and extracellular matrix organization, which rely on endoplasmic reticulum (ER)-localized translation.^47,48^ Conversely, transcripts downregulated in aged ribosomes were enriched in genes encoding proteins involved in translation and spliceosome assembly, which are preferentially translated by free, non-ER-bound ribosomes.^49^ Indeed, upregulated transcripts display enrichment for signal peptides and other signatures of ER-targeted translation (Figure 2M, Figure S3G), whereas downregulated transcripts were depleted of these features. Interestingly, codon frequency analysis reveals that upregulated transcripts encode proteins enriched in hydrophobic residues, whereas downregulated transcripts preferentially encode basic and charged amino acids (Figure 2N). These findings suggest that molecular age-dependent translation is influenced by the physicochemical properties of the nascent polypeptide, possibly through altered interactions of basic residues with the ribosomal exit tunnel that slow down translation elongation,^50^ and may become amplified by additional alterations to codon decoding or elongation dynamics caused by molecular aging.

### Ribosome molecular aging compromises elongation dynamics at basic regions and increases collision propensity

To further probe the mechanistic basis of molecular age-dependent translation, we examined ribosome distributions across regulated transcripts. Ribosome occupancy profiles of translationally downregulated transcripts (TL-DN) revealed a pronounced accumulation immediately downstream of the start codon, followed by a progressive decline across the coding sequence (Figure 3A,B and Figure S4A,C). This pattern is indicative of molecular age-dependent defects in translation elongation dynamics, and suggests an increased incidence of premature translation termination during molecular aging.

**Figure 3.**
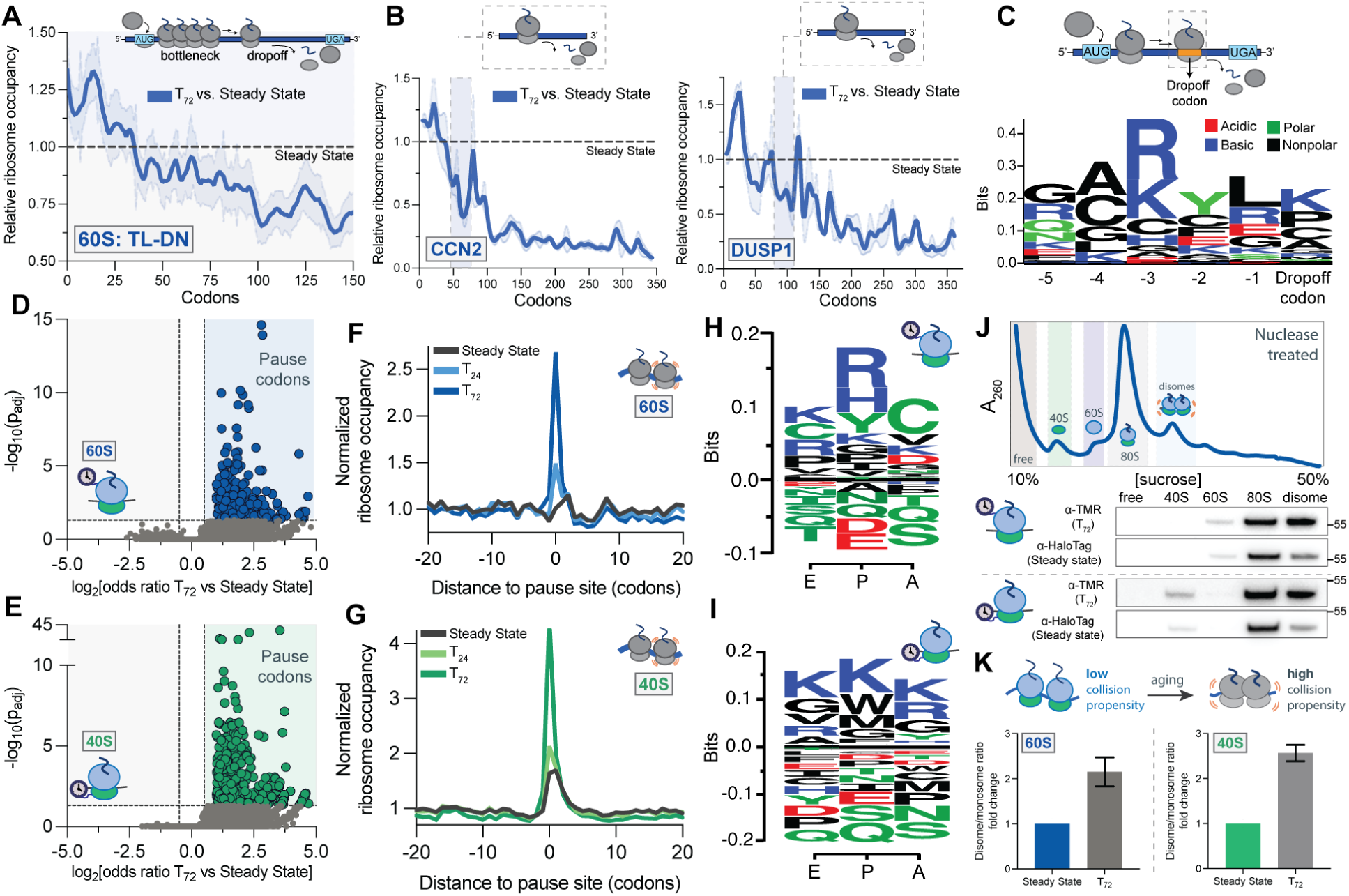
Ribosome molecular aging compromises elongation dynamics at basic regions and increases collision propensity. (A) Metagene analysis of translationally downregulated transcripts (60S, n = 100) illustrating molecular age-associated (T_72_) 60S ribosome elongation bottlenecks and drop-off relative to steady-state. Shaded area along the curve denotes the standard error of the mean (S.E.M.). (B) Ribosome densities of aged ribosomes (T_72_) relative to steady-state for CCN2 and DUSP1. Shaded area along the curve denotes S.E.M. (C) Logo plot of amino acids identified as aged ribosome (T_72_) 10% dropoff codons (90% of steady state) and their preceding codons. (D) Volcano plot of 60S molecular age-dependent pause sites (n = 233, odds ratio > 1, adjusted P_adj_ < 0.05). (E) Volcano plot of 40S molecular age-dependent pause sites (n = 419, odds ratio > 1, adjusted P_adj_ < 0.05). (F) Normalized ribosome occupancies at 60S molecular age-dependent pause sites. (G) Normalized ribosome occupancies at 40S molecular age-dependent pause sites. (H) Logo plot of amino acids being decoded at 60S molecular age-dependent pause sites. (I) Logo plot of amino acids being decoded at 40S molecular age-dependent pause sites. (J) Representative polysome profile and western blot analysis of distribution of molecularly aged and steady-state ribosomes in nuclease-treated sucrose gradients. (K) Quantification of changes in disome-to-monosome ratios relative to steady-state (n = 2).

To identify sequence features associated with premature termination, we modeled ribosome coverage across downregulated transcripts by fitting logistic functions to ‘dropoff codons’, defined as positions at which normalized ribosome occupancy in aged ribosomes fell below 90% of steady-state levels. This analysis revealed significant enrichment of amino acids known to impede elongation dynamics, including proline (P)^51^ and the basic residues lysine (K) and arginine (R),^50^ both at and immediately upstream of these sites (Figure 3C, Figure S4B). These findings are consistent with the elevated abundance of basic amino acids in downregulated relative to upregulated transcripts (Figure 2N), further suggesting that synthesis of proteins enriched for such residues becomes increasingly challenging for molecularly aged ribosomes.

Premature termination can arise from prolonged ribosome pausing, which promotes ribosome collisions and triggers recruitment of quality-control factors that terminate elongation and degrade incomplete nascent polypeptides.^52–56^ We therefore tested whether molecular aging exacerbates ribosome pausing by identifying codons that have a monotonic, significant age-dependent accumulation of ribosomes.^32^ Aging of both 60S and 40S ribosomes increased pausing across hundreds of codons (Figure 3D,E). The extent of pausing at these sites increased progressively with molecular age (Figure 3F,G). Moreover, although molecularly aged 60S and 40S ribosomes paused at overlapping codons, pausing magnitudes at individual codons were largely uncorrelated between subunits (Figure S4D,E). Sites exhibiting increased pausing with molecular age preferentially affected transcripts encoding proteins involved in ribosome biogenesis, translation and protein folding (Figure S4F). Sequence analysis of pause sites revealed positional enrichment of basic residues across multiple ribosomal active-site positions (Figure 3H,I), in line with the composition of identified drop-off codons (Figure 3C). Moreover, ribosome occupancy at polybasic lysine/arginine (K/R) repeats across transcripts showed progressively stronger slowdowns in a molecular age-dependent manner (Figure S4G,H), demonstrating that molecular aging exacerbates ribosome pausing at polybasic regions.

Prolonged translational pausing can promote collisions, which are highly deleterious events that need to be resolved by the ribosome quality control (RQC) pathway.^53,57–59^ We therefore tested whether molecular aging also increases ribosome collision propensity by performing polysome analysis of nuclease-treated lysates, allowing resolution of nuclease-resistant collided ribosomes (disomes) from monosomes (80S) (Figure 3J). This analysis showed a marked increase in the disome-to-monosome ratio in molecularly aged ribosomes relative to steady-state ribosomes (Figure 3K), demonstrating that molecular aging not only increases pausing, but also elevates collision propensity.

### Molecular aging increases the collision propensity of specific ribosome subpopulations

We next sought to identify biochemical features that define molecularly aged ribosomes. Given the structural and chemical complexity of ribosomes, molecular aging could in principle arise from a wide range of alterations, including protein post-translational modifications,^60^ ribosomal RNA (rRNA) modifications,^61^ and chemical damage to both protein and RNA components.^9^ However, approaches that enable comprehensive, high-resolution mapping and experimental perturbation of many of these features remain limited. We therefore focused on rRNA 2’-*O*-methylation and pseudouridylation as a tractable axis of ribosome heterogeneity. These highly conserved^62^ and functionally critical modifications can occur at variable levels,^63^ together account for ∼95% of known rRNA modification events,^64^ and can be quantitatively profiled using established methods such as RiboMethSeq and BID-Seq.^65–67^

We hypothesized that differential rRNA modification patterns may synergize with ribosome molecular aging to increase the susceptibility of specific ribosome subpopulations to functional decline. To test this, we profiled rRNA modification states following molecular age-selective ribosome purifications, coupled with quantitative modification mapping by RiboMethSeq and BID-Seq. We analyzed translating (monosomes) and collided (disomes) ribosomes isolated from nuclease-treated lysates (Figure 4A), as well as unfractionated, non-nuclease-treated lysates representing the total ribosome pool (Figure S5B). This analysis revealed a selective depletion of 40S ribosomal subunits harboring the conserved^68^ pseudouridine Ψ18S-210 within ribosome collisions driven by molecularly aged 60S subunits (Figure 4B,C). Ψ18S-210 is located in helix 10 of the 40S subunit^69^ and is surface-accessible (Figure S5F), suggesting a potential role in modulating local structural integrity or protein-RNA interactions. Notably, this depletion was not observed in translating monosomes or in intact lysates (Figure S5A,C), indicating that the modification loss is specific to aged ribosomes engaged in collisions rather than reflecting a global, age-dependent compositional shift.

**Figure 4.**
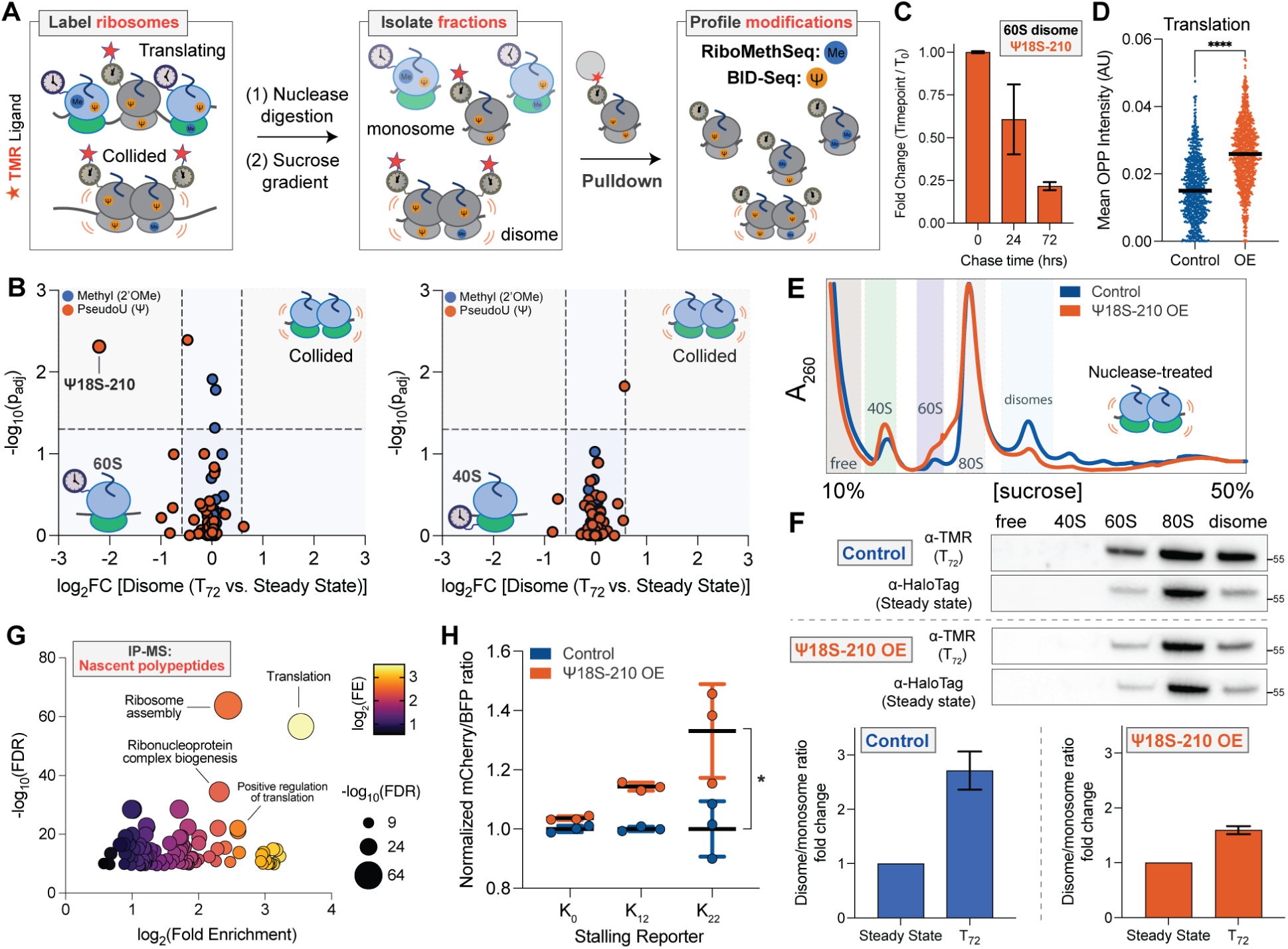
Molecular aging increases the collision propensity of specific ribosome subpopulations. (A) Schematic overview of the experimental strategy used to profile ribosomal RNA modifications in molecular age-dependent translating and collided ribosome subpopulations. (B) Compositional differences of molecularly aged collided 60S and 40S ribosomes (n =2, Fold change (FC) >1.5 fold, p_adj_ < 0.05). (C) Temporal dynamics of Ψ18S-210 in molecular age-dependent 60S ribosome collisions (n = 2). (D) Quantification of newly synthesized polypeptides in Control vs. Ψ18S-210 overexpression cells (n ≥ 848 cells, p < 0.0001, two-tailed Mann-Whitney test). (E) Nuclease-treated polysome analysis of lysates from Control vs. Ψ18S-210 overexpression (OE) cells (F) Quantification of aged and steady-state ribosomes in nuclease-treated sucrose gradients in Control vs. Ψ18S-210 overexpression cells (n = 2). (G) Gene Ontology (Biological Process) terms for nascent polypeptides upregulated by Ψ18S-210 overexpression. (H) Quantification of polybasic stalling reporter readouts in Control vs. Ψ18S-210 overexpression cells (n = 3, p < 0.05, two-tailed Welch’s t test).

We hypothesized that collisions driven by molecularly aged 60S ribosomes require 40S subunits lacking Ψ18S-210. To test this, we depleted the Ψ18S-210(-) subpopulation through snoRNA overexpression (Figure S5D,E). To assess whether depletion of Ψ18S-210(-) ribosomes attenuates molecular age-dependent collisions, we performed polysome profiling following nuclease digestion. In cells overexpressing Ψ18S-210, steady-state disome levels were markedly reduced relative to control cells (Figure 4E), and 60S molecular age-dependent collisions became substantially suppressed (Figure 4F). These findings demonstrate that Ψ18S-210(-) ribosomes are key drivers of 60S molecular age-dependent ribosome collisions.

We next asked whether the observed reduction in collision propensity resulted in measurable changes in total translational output. Nascent polypeptide labeling and polysome profiling showed a marked increase in global translation in the Ψ18S-210 overexpression cells (Figure 4D, Figure S5G). Purification and mass spectrometry analysis of these nascent polypeptides revealed a substantial increase in the production of proteins enriched for basic amino acids, with roles in translation and ribosome assembly, and whose translation decreases as a consequence of molecular aging (Figure 4G; Figure S5J,K). We therefore hypothesized that depletion of Ψ18S-210(-) ribosomes should improve elongation dynamics through polybasic sequences. To test this, we employed dual-fluorescent reporters containing 0, 12, or 22 AAA (K) codons, which induce polybasic-driven ribosome stalling.^70^ Strikingly, Ψ18S-210 overexpression cells exhibited markedly increased reporter readthrough (Figure 4H), demonstrating that Ψ18S-210(-) ribosomes promote polybasic sequence-induced ribosome stalling, with their depletion alleviating elongation defects that are exacerbated by molecular aging.

We hypothesized that alleviation of translational stress through depletion of Ψ18S-210(-) ribosomes would lead to improved cellular fitness. Consistently, growth dynamics were substantially enhanced in Ψ18S-210 overexpression cells compared with control cells (Figure S5H). Furthermore, we tested whether the observed reduction in collision propensity would confer protection from ribotoxic stress. Challenging cells with the collision-inducing drug anisomycin (ANS)^71^ indeed showed that Ψ18S-210 overexpression increased resistance to ribotoxic stress (Figure S5I). Together, these findings identify Ψ18S-210(-) ribosomes as a molecularly vulnerable ribosome subpopulation that becomes collision-prone in concert with molecular aging of 60S ribosomes. Intriguingly, the snoRNA machinery responsible for installing Ψ18S-210 is downregulated in aged human cell cohorts compared to young cells (Figure S5L), suggesting a potential link between this ribosome subpopulation and age-associated proteostasis decline.

### Enriching the pool of aged ribosomes amplifies molecular age-dependent defects in the steady-state translatome

In light of our findings, we hypothesized that imbalances in age demographics of ribosomes, such as those arising under conditions of impaired ribosome biogenesis and turnover, are sufficient to amplify molecular age-dependent defects in the steady-state translatome. To test this, we developed a strategy to selectively target newly synthesized ribosomes for degradation while preserving pre-existing ribosomes, thus enriching aged ribosomes in cells over time (Figure 5A). Pre-existing 60S ribosomes were first saturated with the HaloTag-TMR ligand, preventing subsequent labeling. Cells were then continuously treated with HaloPROTAC3 (PROTAC) or its inactive enantiomer, *ent*-HaloPROTAC3 (Control),^72^ such that newly synthesized ribosomes are selectively labeled with either ligand. This approach led to robust degradation of newly synthesized HaloTag fusions while preserving the older, pre-existing ones (Figure S6A,B). Time-resolved imaging and quantitative analyses demonstrated that PROTAC treatment markedly reduced ribosome turnover (Figure 5B,C) by diminishing both the dilution and degradation components (Figure 5D), thereby enriching the population of aged ribosomes within cells.

**Figure 5.**
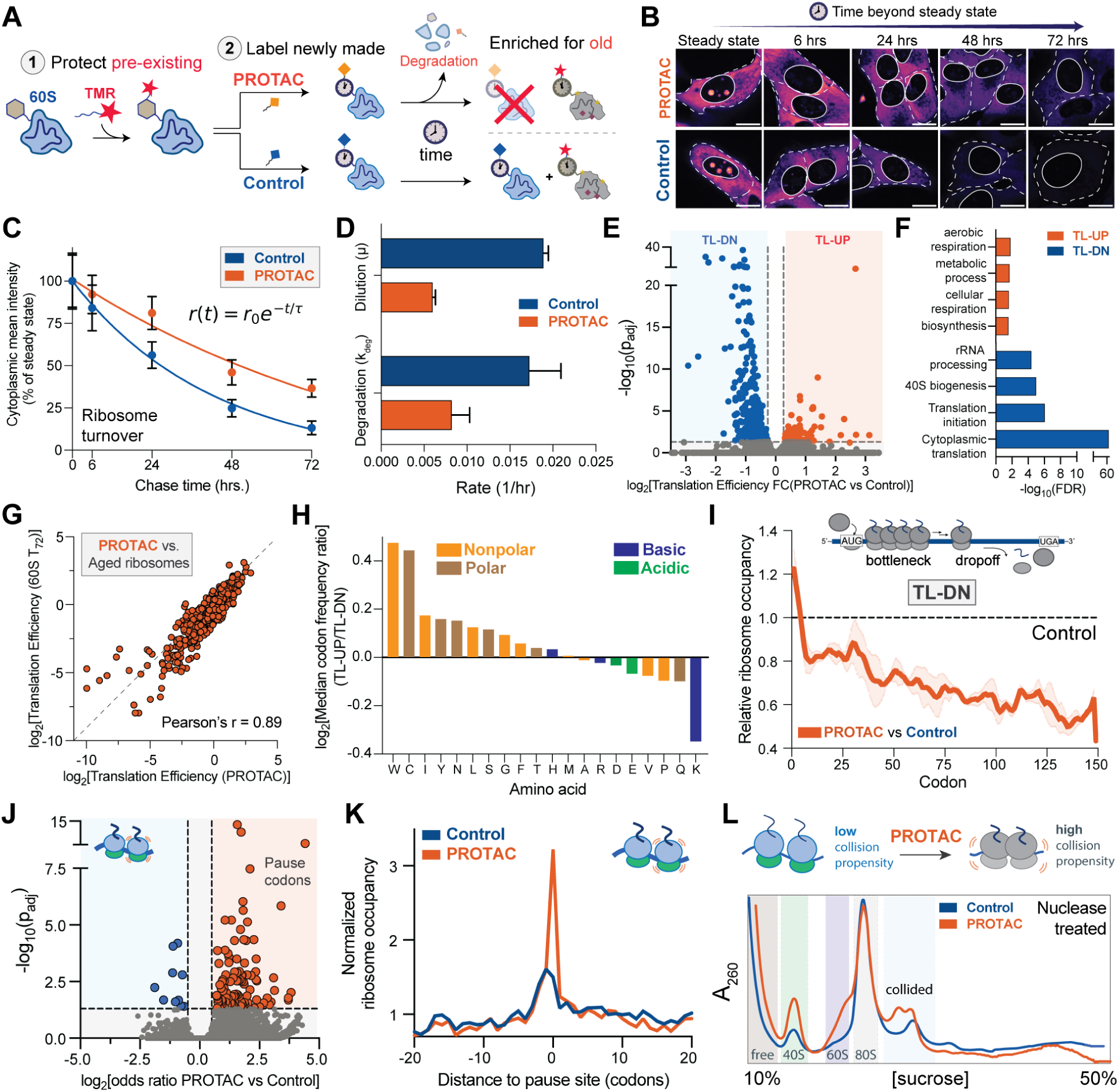
Enriching the pool of aged ribosomes amplifies molecular age-dependent defects in the steady-state translatome. (A) Schematic overview of the experimental strategy used to enrich aged ribosome populations in cells. (B) Time-resolved imaging (Scale bar = 15 μm) and (C) quantification of ribosome turnover dynamics in control versus PROTAC-treated cells (mean ± S.E.M., n ≥ 38 cells). (D) Quantification of ribosome dilution and degradation rates under Control and PROTAC treatment (mean ± S.E.M., n ≥ 38 cells). (E) Volcano plot of translation efficiency changes between control and PROTAC-treated cells (n = 529 regulated transcripts, TL-DN = 414, TL-UP = 115, adjusted p<0.05, FC > 1.2 fold). (F) Gene Ontology analysis (Biological process) for translationally upregulated (TL-UP) and translationally downregulated (TL-DN) transcripts under PROTAC relative to Control. (G) Correlation between translation efficiencies in PROTAC-treated cells and aged 60S ribosomes (T_72_) at PROTAC-regulated transcripts. (H) Ratio of median codon frequencies between TL-UP and TL-DN transcripts regulated by PROTAC condition. (I) Metagene analysis of translationally downregulated transcripts in PROTAC condition (mean ± S.E.M., n = 100) illustrating elongation defects relative to control. (J) Volcano plot of PROTAC-dependent pause sites (increased pausing n = 108, odds ratio > 1, adjusted P < 0.05). (K) Normalized ribosome occupancies at PROTAC-dependent pause sites. (L) Nuclease-treated sucrose gradients in control and PROTAC-treated conditions.

We next asked whether molecular age-dependent translational defects became amplified at steady state under this PROTAC treatment. Ribo-seq revealed extensive remodeling of translation efficiencies across hundreds of transcripts in PROTAC- versus Control-treated cells (Figure 5E,F, Figure S6E-G,J). Strikingly, translation efficiencies of PROTAC-regulated transcripts were highly correlated with those observed for molecularly aged 60S ribosomes (T_72_) (Pearson’s r = 0.89; Figure 5G), demonstrating that shifts in ribosome age demographics are sufficient to amplify molecular age-dependent translational programs in cells.

Analysis of regulated transcripts revealed compositional preferences that closely mirrored those observed in molecularly aged 60S ribosomes (Pearson’s r = 0.79, Figure 5H, Figure S6H). Translationally upregulated transcripts were enriched in non-polar amino acids, whereas downregulated transcripts showed a relative enrichment in basic residues, particularly lysine. Because molecular aging impairs translation elongation, we examined whether PROTAC treatment similarly disrupted elongation dynamics. PROTAC treatment recapitulated elongation defects characteristic of molecularly aged ribosomes, including start codon-proximal bottlenecks and premature termination across translationally downregulated transcripts (Figure 5I). In addition, PROTAC treatment caused a substantial increase in ribosome pausing (Figure 5J,K), predominantly affecting transcripts involved in cytoplasmic translation (Figure S6I). Sequence analysis revealed positional enrichment of basic residues across multiple ribosomal active-site positions (Figure S6K) and dose-dependent effects at polybasic stretches of K/R repeats (Figure S6L), indicating that imbalances in ribosome age demographics are sufficient to compromise translation elongation at regions rich in basic amino acids. Consistent with these defects, sucrose gradient fractionation of nuclease-treated lysates showed that PROTAC treatment amplified ribosome collisions at steady state (Figure 5L), mirroring the collision-prone state of molecularly aged ribosomes.

Together, these results demonstrate that shifting the ribosome pool towards older ribosomes is sufficient to amplify molecular age-dependent translation defects and suggests that organismal aging and disease-associated perturbations in the biogenesis and turnover of ribosomes could similarly enrich aged ribosomes, contributing to functional failures such as proteostasis collapse.^32,34,73^

### Ribosome molecular aging shapes translation dynamics during organismal aging

Given the translational defects associated with molecularly aged ribosomes, we next asked whether there was a link between molecular aging and organismal aging. To directly track aged ribosomes during organismal aging, we generated a *C. elegans* strain in which the endogenous *rpl10a* locus was fused to HaloTag (Figure S7A). HaloTag ribosomes in this strain retained functional activity and were properly assembled into actively translating pools (Figure S7B). We optimized ligand labeling and chase conditions to enable temporal marking of aged ribosome demographics (Figure S7C,D). In this experiment, Day 1 adult worms were pulse labeled with the HaloTag-TMR ligand for 24 hours to label pre-existing ribosomes, followed by extensive washout at Day 2 to remove residual ligand. Animals were then aged without further labeling until Day 10 of adulthood (Figure 6A). Under these conditions, TMR ligand-labeled ribosomes represented a population that had persisted for at least 8 days *in vivo*, whereas the total HaloTag signal reported the complete pool of all tagged ribosomes regardless of their molecular age.

**Figure 6.**
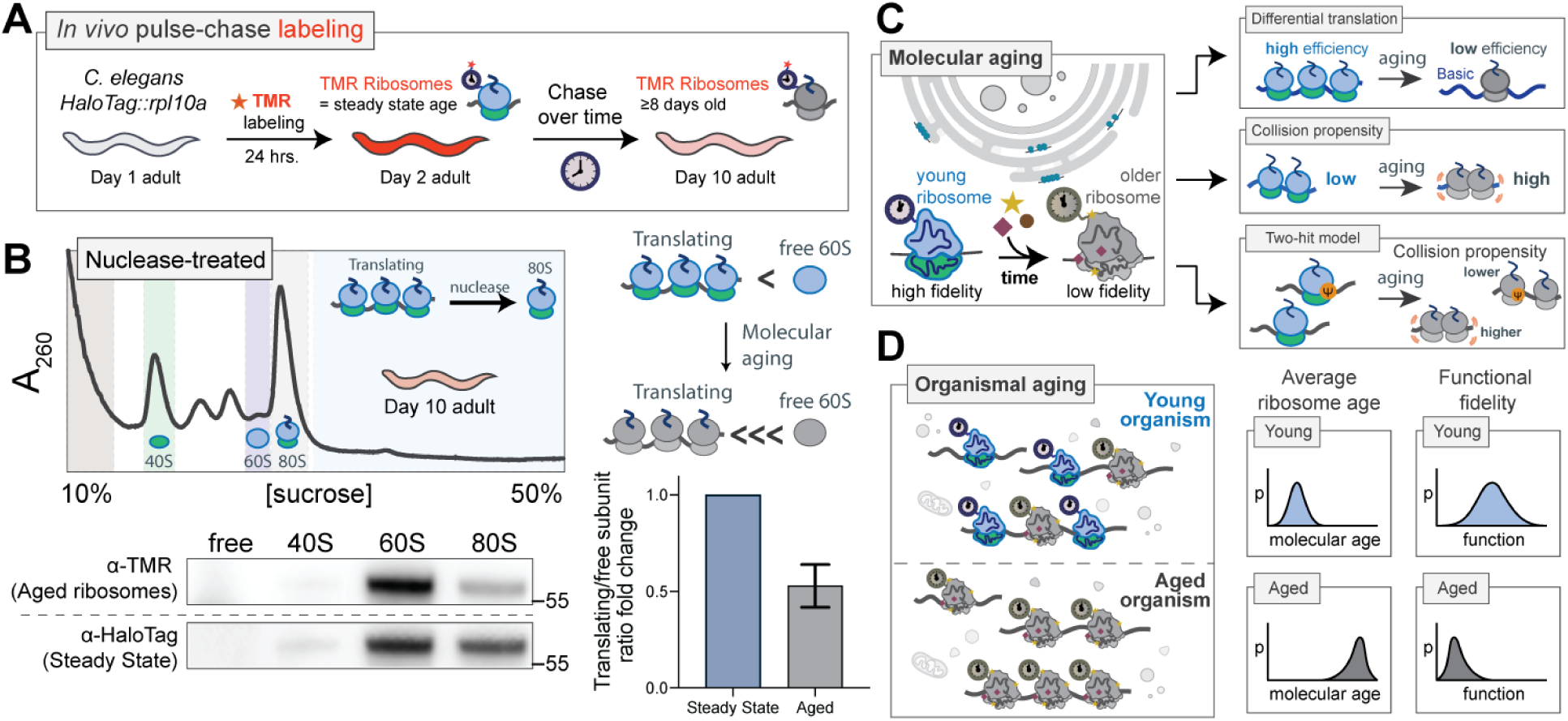
Ribosome molecular aging shapes translation dynamics during organismal aging. (A) Schematic of the experimental strategy for *in vivo* labeling of aged ribosomes in the *HaloTag::rpl10a C. elegans* strain. (B) Polysome analysis of nuclease-treated lysates from pulse-chase-labeled *C. elegans* reveals molecular age-dependent shifts in distribution between free and actively translating ribosomal subunit pools (n = 2). (C) Model of how molecular aging changes ribosome composition and interactions, shifting ribosomes from high- to lower-fidelity functional states over time. These changes influence the translation efficiency of mRNAs rich in basic amino acids, and overall increase ribosome collision propensity. A two-hit model is proposed in which specific ribosome subpopulations become collision prone through the molecular aging process. (D) Schematic showing how at the organismal level, physiological aging raises the average ribosome age, shifting the distribution toward lower functional fidelity.

To compare the translational status of aged ribosomes with that of the total ribosome pool in aged animals, we performed sucrose gradient fractionation of lysates from aged *C. elegans* coupled with nuclease treatment to collapse actively translating polysomes into 80S monosomes. Gradient fractions were analyzed by western blot using probes against the TMR ligand to detect aged ribosomes and against HaloTag protein to detect the steady state pool of tagged ribosomes. This approach thus enabled parallel measurement of the distribution of aged and total ribosomes across free subunits and translating 80S fractions. The ratio of translating (80S) ribosomes to free subunits (60S) was substantially lower in aged ribosomes compared to the total pool (Figure 6B). These results indicate that molecularly aged ribosomes *in vivo* display reduced translational activity relative to the overall ribosome population at the same organismal age, consistent with molecular age-dependent alterations that compromise their function.

Together, these data demonstrate that ribosomes that have undergone extended residence *in vivo* show distinct translation dynamics. This age-dependent shift in ribosome translational status supports a model in which molecular aging of ribosomes contributes directly to altered translation dynamics during organismal aging (Figure 6C,D).

## DISCUSSION

The long-lived nature of many cellular components, including ribosomes, has been recognized for decades. However, whether their prolonged stability contributes meaningfully to their functional landscape has remained unresolved, largely owing to the lack of tools to interrogate molecular age-dependent functional states in living cells. Here we have developed and applied a method to map distinct stages of the ribosome life cycle across short and extended molecular timescales, revealing previously uncharacterized functional trajectories that emerge long after maturation. These findings establish ribosome molecular age as an intrinsic axis of functional heterogeneity, with direct consequences for the dynamics of translation.

Molecular aging of ribosomes drives differential translation of distinct mRNA subsets defined by the physical properties of their nascent polypeptides, demonstrating that the translational status of an mRNA is not an intrinsic property of the sequence, but also a property of the molecular age of the ribosome decoding it. Mechanistically, molecular aging increases vulnerability to coding regions rich in basic amino acids, impairing elongation and promoting ribosome pausing, collisions, and premature termination. As a result, molecular aging selectively hinders the synthesis of proteins enriched in basic residues, with important consequences for proteostasis and ribosome quality control.

Manipulation of ribosome age demographics demonstrated that aging of the ribosome pool is sufficient to recapitulate molecular age-dependent translational defects. A PROTAC-based strategy that biased the population toward older ribosomes amplified elongation defects across the translatome and recapitulated elevated ribosome pausing, premature termination, and collision propensity at steady state. Ribosome degradation rates decreased following this perturbation, suggesting the existence of cellular mechanisms that dynamically tune ribosome stability in response to production rates, which could be elucidated in the future through functional screening approaches.^74,75^

Our findings indicate that faithful translation of distinct mRNA subsets, especially those encoding basic amino acid-rich proteins such as components of the translation machinery, requires sustained ribosome biogenesis flux. Molecular aging may therefore function in part to couple a biosynthetic capacity to functional need, progressively limiting processes like translation and ribosome biogenesis under growth-limiting conditions. This framework is interesting in the context of the persistent requirement for ribosome synthesis in post-mitotic cells that do not experience ribosome dilution through cell division, such as neurons and myocytes,^76,77^ and supports a model in which specialized turnover pathways selectively eliminate deleterious ribosome species to maintain long-term proteostasis.

We show that molecular aging increases the collision propensity of specific ribosome subpopulations, indicating differential susceptibility among ribosome pools to molecular age-associated decline. Notably, susceptible ribosome subpopulations lacked a particular pseudouridine (Ψ18S-210), a modification that could serve to protect rRNA from hydrolysis or RNase-mediated degradation.^78^ Together, these results support a “two-hit” model in which specifically susceptible ribosomes become functionally deleterious with increasing molecular age (Figure 6C). Using snoRNA overexpression, depletion of the susceptible Ψ18S-210 ribosome subpopulation ameliorated functional defects observed upon molecular aging, such as polybasic stalling and ribosome collisions, and enhanced resistance to ribotoxic stress. Thus, selectively targeting this or other specific ribosome subpopulations may represent a therapeutic strategy for diseases characterized by imbalances in ribosome age demographics, including age-related disorders. Indeed, we observed that the snoRNA machinery responsible for installing Ψ18S-210 is downregulated in aged human cell cohorts relative to young cells (Figure S5L), supporting further investigation into the role of this ribosome subpopulation in aging-associated proteostasis defects, and other opportunities for therapeutic intervention.

Defining the precise molecular features that distinguish ribosome age demographics and drive age-dependent functional dynamics will require further investigation, but likely include time-dependent protein composition^79^ and post-translational modifications,^60^ or stochastic chemical damage such as oxidation^80,81^ or hydrolysis^82^. Such features may accumulate in the absence of dedicated exhaustive ribosome repair pathways, which have been described for a limited set of ribosomal components,^9^ in contrast to cellular components like DNA.^83^

Interestingly, our data point to the accumulation of damage in both aged small subunits (40S), and large subunits (60S) as contributing to translational dysregulation. Moreover, there is clear cross talk between the physiochemical state of the two subunits, as molecularly aged 60S are particularly vulnerable when assembled into ribosomes with 40S subunits lacking the Ψ18S-210 modification. Future work will be required to mechanistically dissect this coupling.

Recent studies have reported translation defects that intensify with organismal aging in systems such as *C. elegans*, *S. cerevisiae,* and the African turquoise killifish.^32,34^ In these models, organismal aging exacerbates ribosome pausing at regions rich in basic amino acids, promoting ribosome collisions and driving the accumulation of incomplete nascent polypeptides that overwhelm proteostasis networks and promote pathological protein aggregation. These previous findings on organismal aging mirror those we report here, on molecular age-dependent decline in ribosome function. This suggests that molecular aging of ribosomes and other components of the translational machinery could directly drive these organismal aging phenotypes, which are indeed associated with declines in ribosome biogenesis and turnover.^29,30,84,85^

Consistent with this concept, our findings in aged *C. elegans* provide the first direct *in vivo* evidence that long-lived ribosomes display altered translational activity compared to the steady-state ribosome population. Thus, ribosome molecular aging contributes to altered translation dynamics during organismal aging, although the extent to which the underlying mechanisms operating in these aged, post-mitotic organisms parallel those occurring in human cells remains to be determined. Future work in *C. elegans* and other model organisms using higher-resolution approaches, such as molecular age-selective Ribo-seq will be necessary to further dissect how proteostasis decline during organismal aging may be primarily driven by aged ribosomes.

Molecular aging likely causes time-dependent functional changes beyond ribosomes, representing a general property of long-lived cellular components. As with non-living materials that undergo progressive physicochemical changes over time, age-associated molecular deterioration may gradually alter the functional properties of these components, thereby shaping outcomes across diverse biological processes ranging from regulation of central dogma processes to maintenance of cellular structure. The framework described here establishes a general method for tracking the functional performance of macromolecular complexes over time, enabling future studies to uncover molecular age-dependent functional changes across other persistent cellular components, informing therapeutic avenues for aging-related disease.

## LIMITATIONS OF THE STUDY

Several important questions remain unresolved. The precise chemical alterations that drive ribosome molecular aging are still unknown. Although we identify ribosomal RNA modifications that are altered in molecular age-dependent ribosome collisions, how these changes interact with other age-dependent chemical alterations and contribute mechanistically to functional decline remains unclear. While our approach resolves distinct ribosome demographics across space and time, their dynamic behavior in living cells has not yet been fully measured and may benefit from single-molecule approaches.^87–91^ Our method relies on HaloTag labeling, which is highly efficient but may not be broadly applicable for endogenous tagging of some cellular components due to size limitations; further development of compact, efficient self-labeling tags may be necessary to expand this strategy to smaller cellular constituents. Our PROTAC strategy may impact the translatome through mechanisms besides enrichment of aged ribosomes. Finally, although we demonstrate that molecularly aged ribosomes in aged *C. elegans* exhibit altered functional activity compared to the steady-state pool, it remains unknown whether molecular aging affects *C. elegans* ribosomes through mechanisms analogous to those observed in human cells. Addressing this question in detail will require transcriptome-wide translational profiling approaches, such as Ribo-seq, to define the global impact of ribosome aging on translational output.

## RESOURCE AVAILABILITY

All materials and resource requests should be directed to and will be fulfilled by the lead contact, Clifford P. Brangwynne.

## DATA AND CODE AVAILABILITY

Data in this manuscript will be shared by the lead contact upon request. Data supporting these findings will be made available on public databases (GEO, ProteomeXchange) upon publication or upon request. Custom scripts used for this study are provided in the following repository: https://github.com/SoftLivingMatter/Botello-2026-ribosome-aging.

## ACKNOWLEDGEMENTS

We are grateful to Audrey Zhu and Miguel Iglesias of the Wuhr laboratory for assistance with sucrose gradient fractionation; Evangelos G. Gatzogiannis for microscopy assistance; Sofia A. Quinodoz for help with sequencing library preparation; Aaron Lin for help with nucleofections; Christina DeCoste for assistance with cell sorting; Nima Jaberi-Lashkari and Nikhil Patel for discussions and feedback; David W. Sanders for reagents and discussions, and all Brangwynne lab members for helpful discussions as well as experimental support. This work was supported by the Howard Hughes Medical Institute (HHMI), the Chan Zuckerberg Initiative Exploratory Cell Network, the St. Jude Research Collaborative on the Biology and Biophysics of RNP granules, and the Princeton Center for Complex Materials (DMR-2011750). I.M.C. is supported by Paul G. Allen Family Foundation and the NIH NIAID (R01AI174515). P.J.M. is supported by an NIH F31 fellowship (G0001-10017556-101). J.F.B. and L.W.W. are supported by an NSF graduate research fellowship. The work by Q.Y. was supported in part by a Harold W. Dodds Fellowship from Princeton University.

## AUTHOR CONTRIBUTIONS

J.F.B. and C.P.B. designed the study. J.F.B, L.J., P.J.M., A.A.A., M.S.E., M.L., L.W.W, and C.J.W. performed experiments. J.F.B., L.J., P.J.M., A.A.A., M.B. and T.J.C. performed imaging, genomics and proteomics analysis. Q.Y., C.P.B., and A.K. performed modeling. P.J.M. worked under the supervision of I.M.C. J.F.B. and C.P.B. wrote the manuscript with input from all authors. J.F.B. made the figures with contributions from all authors.

## DECLARATION OF GENERATIVE AI AND AI-ASSISTED TECHNOLOGIES

During the preparation of this manuscript, ChatGPT was used to enhance the readability and clarity of certain sections. All content generated with the assistance of this tool was reviewed and edited by the authors, who take full responsibility for the final version of the publication.

## SUPPLEMENTAL FIGURES

**Figure S1.**
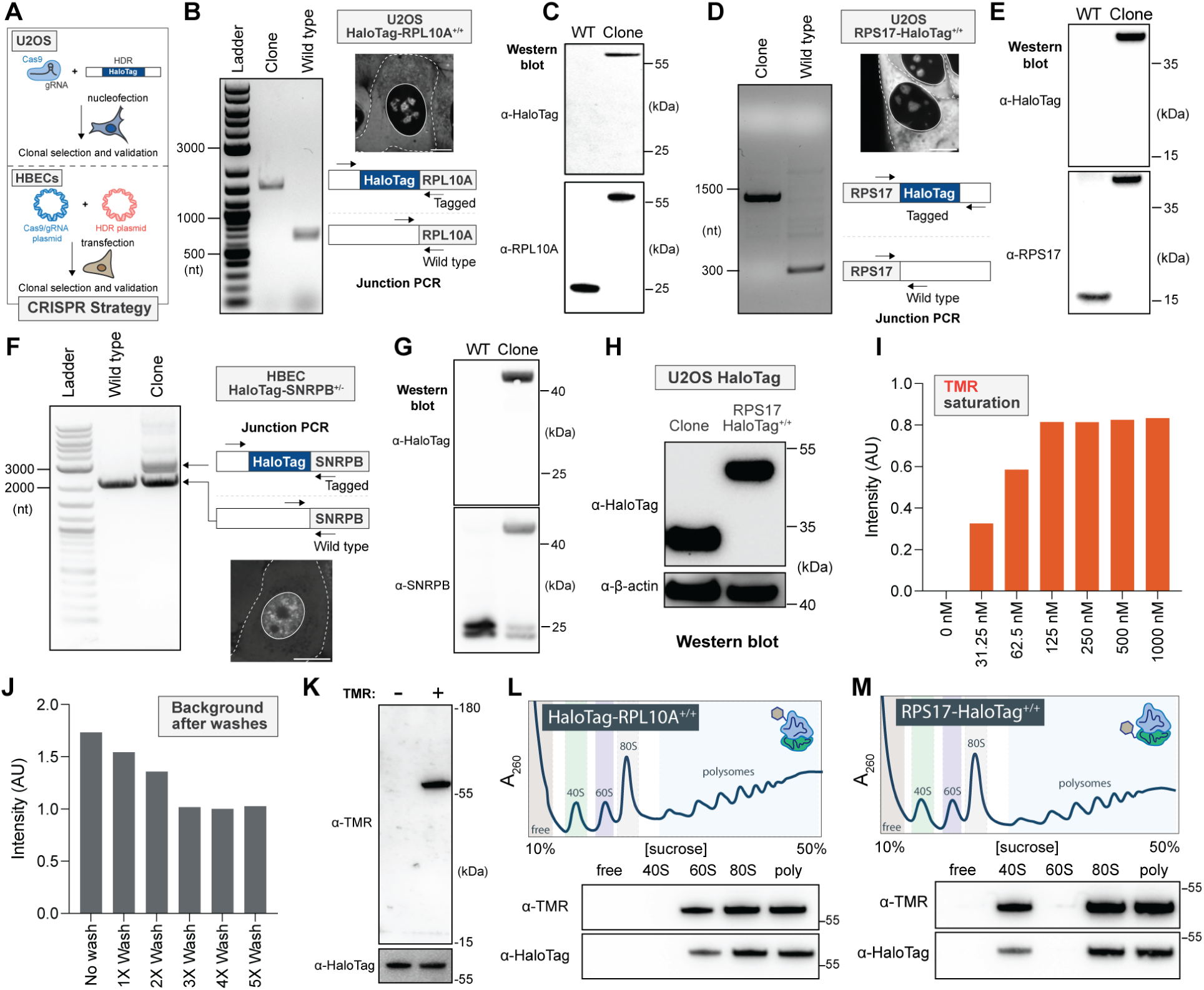
Cell line engineering, validation, and HaloTag-TMR labeling. (A) Strategy for tagging proteins of interest using CRISPR/Cas9 in U2OS cells and HBECs. (B) Genomic junction PCR demonstrating homozygous tagging of RPL10A with HaloTag on its N-terminus and image of the resulting clone showing resulting localization in U2OS (Scale bar = 10 μm). (C) Western blot analysis of HaloTag and RPL10A in the resulting clone compared to wild type U2OS cells. (D) Genomic junction PCR demonstrating homozygous tagging of RPS17 with HaloTag on its C-terminus and image of the resulting clone showing resulting localization in U2OS (Scale bar = 10 μm). (E) Western blot analysis of HaloTag and RPS17 in the resulting clone compared to wild type U2OS cells. (F) Genomic junction PCR demonstrating heterozygous tagging of SNRPB with HaloTag on its N-terminus and image of the resulting clone showing resulting localization in HBECs (Scale bar = 10 μm). (G) Western blot analysis of HaloTag and SNRPB in the resulting clone compared to wild type HBECs. (H) Western blot analysis of HaloTag expressed by lentiviral transduction in wild-type U2OS cells demonstrating expression comparable to ribosomal proteins of interest. (I) Saturation curve of HaloTag-TMR labeling in U2OS RPS17-HaloTag^+/+^ cells (n ≥ 30 cells). (J) HaloTag-TMR background levels after incubation, followed by multiple washes in wild-type U2OS cells (n ≥ 55 cells). (K) Extended view of blot in Figure 1B ; western blot analysis demonstrating specific recognition of HaloTag-TMR-labeled HaloTag-RPL10A^+/+^ by TRITC polyclonal antibody. (L,M) Polysome analysis of endogenously tagged HaloTag-RPL10A^+/+^ and RPS17-HaloTag^+/+^ clones, respectively, demonstrating effective HaloTag incorporation and TMR labeling across ribosome fractions.

**Figure S2.**
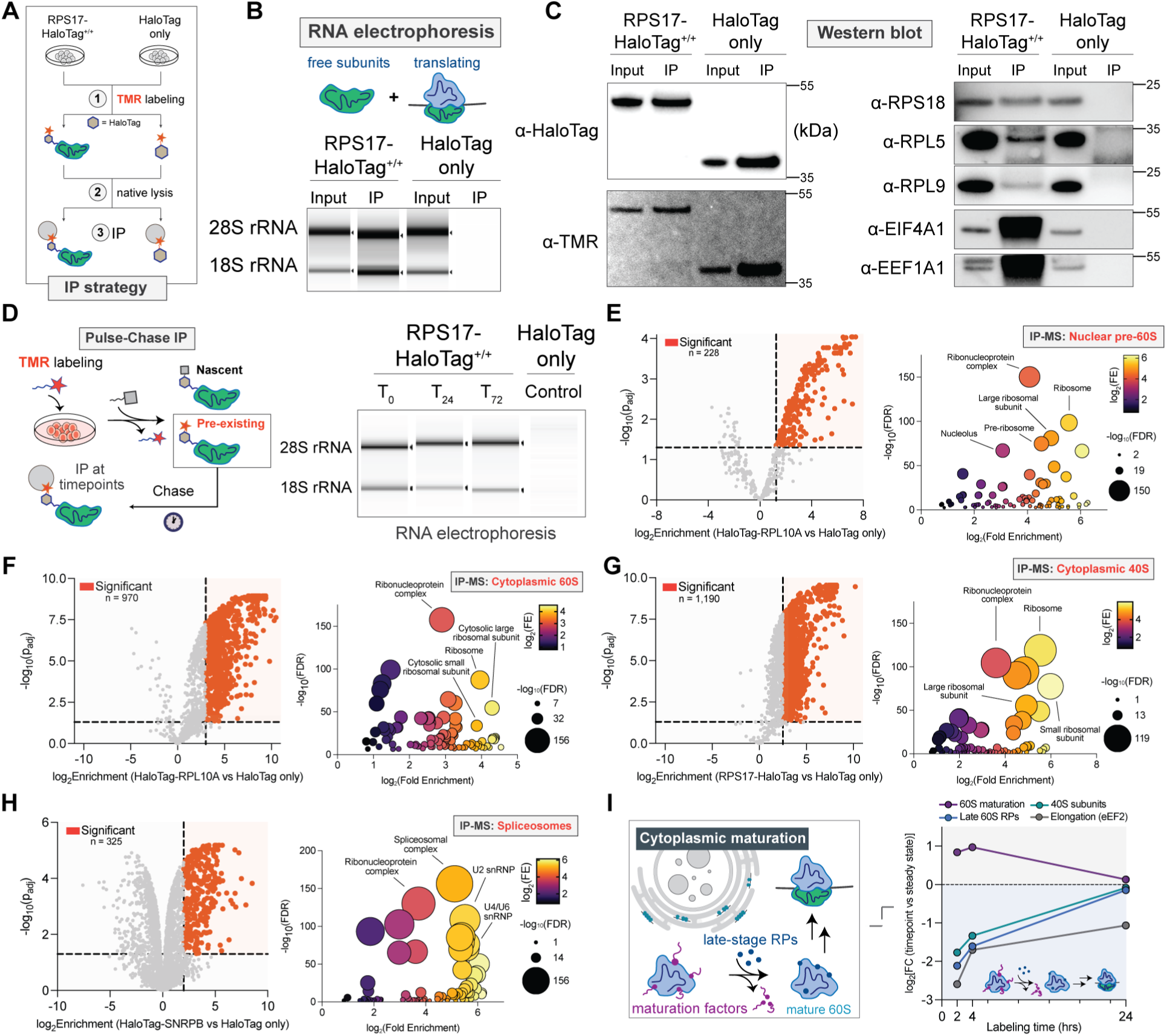
Native immunopurification of cellular complexes through HaloTag-TMR labeling. (A) Strategy for native immunopurification (IP) of small ribosomal subunits via HaloTag-TMR labeling. (B) RNA electrophoresis of input and IP elution products from RPS17-HaloTag^+/+^ and HaloTag-only control U2OS cells. (C) Western blot analysis of input and IP elution products from RPS17-HaloTag^+/+^ and HaloTag-only control cells demonstrating purification of various ribosome components in an RPS17-HaloTag-dependent manner. (D) Pulse-chase IP strategy in RPS17-HaloTag^+/+^ cells demonstrating ribosome IP across extended molecular timescales following chase (0, 24, and 72 hrs). (E) Volcano plot and Gene Ontology (cellular component) analysis of immunopurification-mass spectrometry (IP-MS) data from nuclear pre-60S subunits purified from HaloTag-RPL10A^+/+^ U2OS cells (log_2_FC > 1.25, p_adj_ < 0.05, n = 228 proteins, from n = 3 biological replicates). (F) Volcano plot and Gene Ontology (cellular component) analysis of IP-MS data from cytoplasmic 60S subunits purified from HaloTag-RPL10A^+/+^ U2OS cells (log_2_FC > 3, p_adj_ < 0.05, n = 970 proteins, from n = 3 biological replicates). (G) Volcano plot and Gene Ontology (cellular component) analysis of IP-MS data from cytoplasmic 40S subunits purified from RPS17-HaloTag^+/+^ U2OS cells (log_2_FC > 2.5, p_adj_ < 0.05, n = 1,190 proteins, from n = 3 biological replicates). (H) Volcano plot and Gene Ontology (cellular component) analysis of IP-MS data from spliceosomes purified from HaloTag-SNRPB^+/-^ HBEC cells (log_2_FC > 2, p_adj_ < 0.05, n = 326 proteins, from n = 3 biological replicates). (I) Dynamics of cytoplasmic nascent 60S interactions quantified by IP-MS of time-resolved intermediates (n = 3).

**Figure S3.**
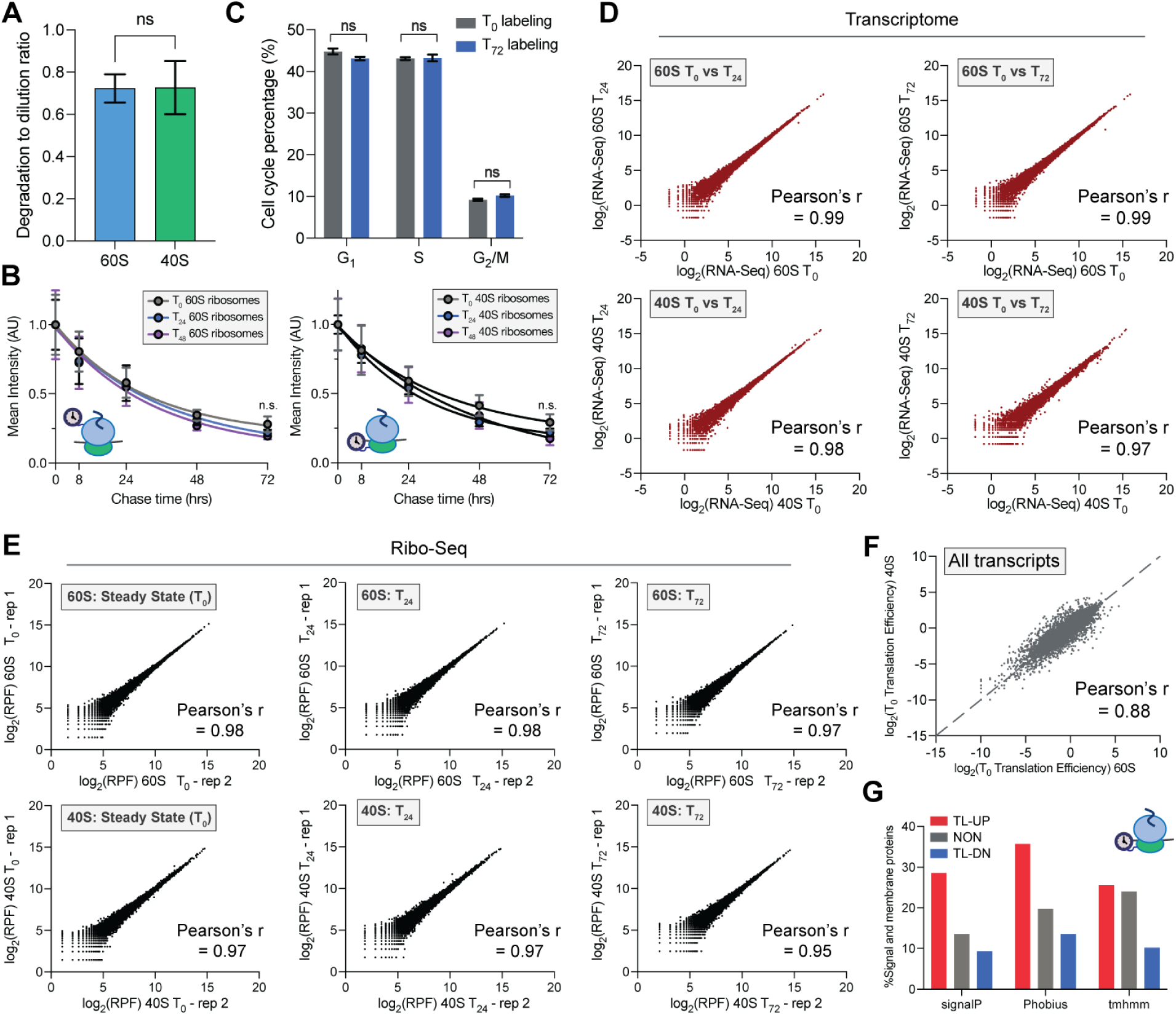
Characterization of ribosome turnover and molecular age-dependent Ribo-seq experiments. (A) Comparative analysis of degradation-to-dilution ratio differences between 60S and 40S ribosomes (n ≥ 57 cells, p = ns, two-tailed Welch’s test). (B) Comparative analysis of turnover rates for 60S and 40S ribosomes of different ages, where T_0_ is steady-state ribosomes and T_24_ and T_48_ refer to ribosomes at least 24 and 48 hr old, respectively (n ≥ 178 cells, p > 0.05, two-tailed Mann-Whitney test). (C) Cell cycle distributions under steady-state (T_0_) and long-term ribosome labeling conditions (T_72_) (n = 3, p = ns, two-tailed Welch’s test). (D) Comparative transcriptome correlation analysis across multiple conditions in molecular age-dependent Ribo-seq experiments. (E) Correlations between replicates for molecular age-dependent Ribo-seq experiment (RPF = Ribosome-protected footprint). (F) Correlation between steady state (T_0_) translation efficiencies of 60S and 40S Ribo-seq experiments. (G) Percent of signal and membrane proteins encoded by transcripts regulated by aged 40S ribosomes according to multiple algorithms.

**Figure S4.**
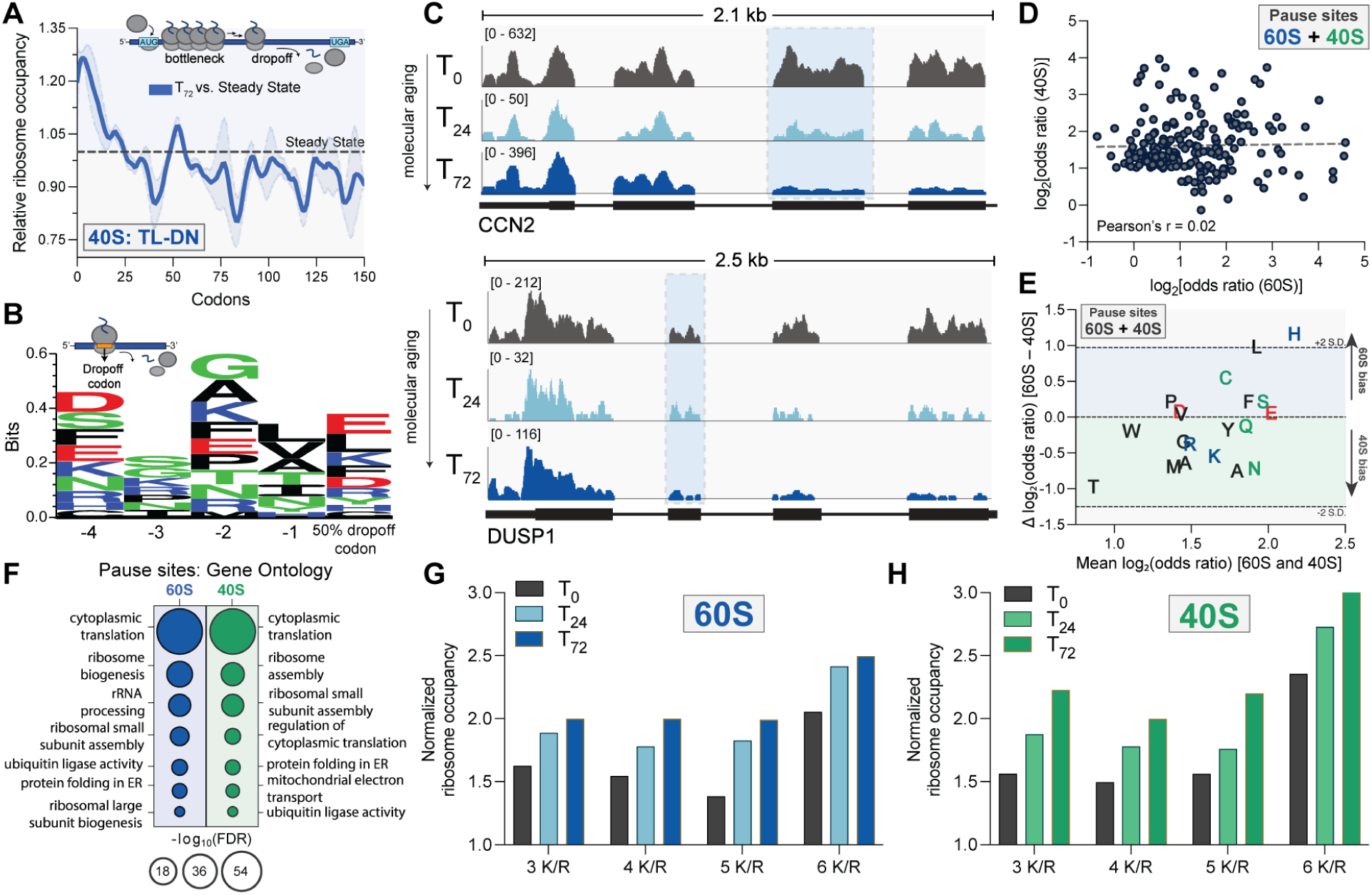
Additional characterization of ribosome molecular age-dependent elongation dynamics. (A) Metagene analysis of translationally downregulated transcripts illustrating molecular age-associated (T_72_) 40S ribosome elongation bottlenecks and drop-off relative to steady-state. (B) Logo plot of amino acids identified as aged ribosome (T_72_) 50% dropoff codons and their preceding codons. (C) Transcript-wide view of molecular age-dependent ribosome occupancy for genes represented in Figure 3B. (D) Correlation between pausing magnitudes for the union of 60S and 40S molecular age-dependent pause sites (E) Bland-Altman analysis of amino acid-dependent biases in pause strengths at 60S and 40S sites. (F) Gene Ontology analysis (Biological Process) of genes with 60S or 40S molecular age-dependent pause sites. (G) Average ribosome occupancy at polybasic regions of different lengths for steady state (T_0_) or molecularly aged (T_24_, T_72_) 60S ribosomes. (H) Average ribosome occupancy at polybasic regions of different lengths for steady state (T_0_) or molecularly aged (T_24_, T_72_) 40S ribosomes.

**Figure S5.**
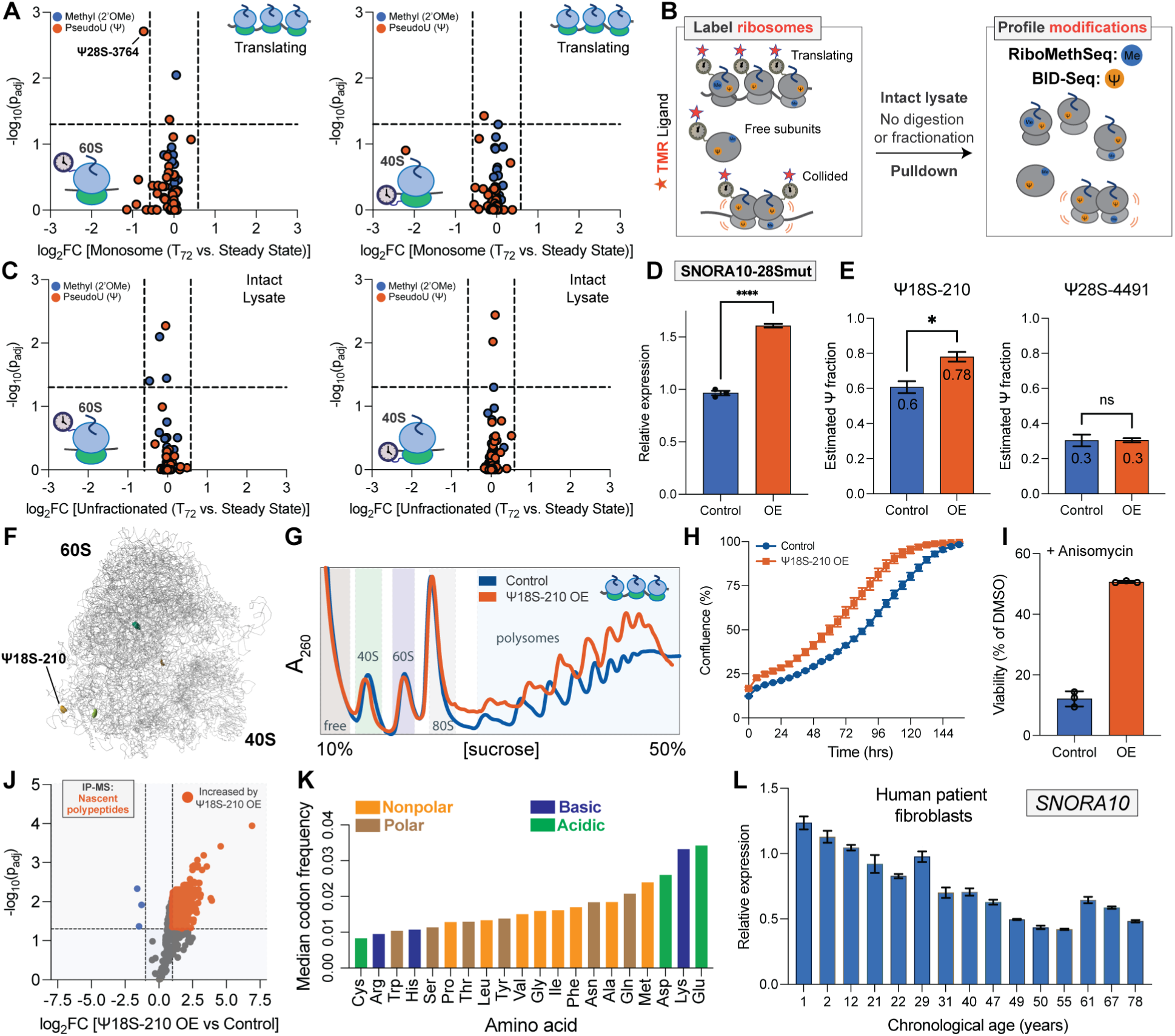
Additional characterization of molecular age-dependent ribosomal RNA modifications and their impact on cellular phenotypes. (A) Compositional differences of molecularly aged translating (monosome) 60S and 40S ribosomes (n = 2, Fold change (FC) >1.5 fold, p_adj_ < 0.05). (B) Schematic overview of the experimental strategy used to profile molecular age-dependent ribosomal RNA modifications in intact, unfractionated lysates. (C) Compositional differences of molecularly-aged 60S- and 40S-associated ribosomes (Fold change (FC) >1.5 fold, p_adj_ < 0.05). (D) Quantitative PCR analysis of SNORA10-28Smut (lacking Ψ28S-4491 modification activity) overexpression relative to control cells (n = 3, p < 0.0001, two-tailed Welch’s t test). (E) BIHIND-qPCR analysis of ribosomal RNA modification overexpression in total RNA (Ψ18S-210: p < 0.05, Ψ28S-4491: p = ns ; two-tailed Welch’s t test). (F) Location of Ψ18S-210 in the structure of the ribosome (PDB: 6QZP). (G) Polysome analysis comparing control and Ψ18S-210 overexpression cells. (H) Growth dynamics of control and Ψ18S-210 overexpression cells (n = 5). (I) Resistance of Ψ18S-210 overexpression (OE) cells to anisomycin treatment relative to control. (J) Volcano plot of IP-MS data of purified nascent polypeptides from Control vs. Ψ18S-210 overexpression (OE) cells (log_2_FC > 1, p_adj_ < 0.05, n = 765 upregulated proteins from n = 3 biological replicates). (K) Median codon frequencies of upregulated nascent polypeptides. (L) Quantitative PCR analysis of SNORA10 expression in human patient fibroblasts of increasing chronological age (n = 3).

**Figure S6.**
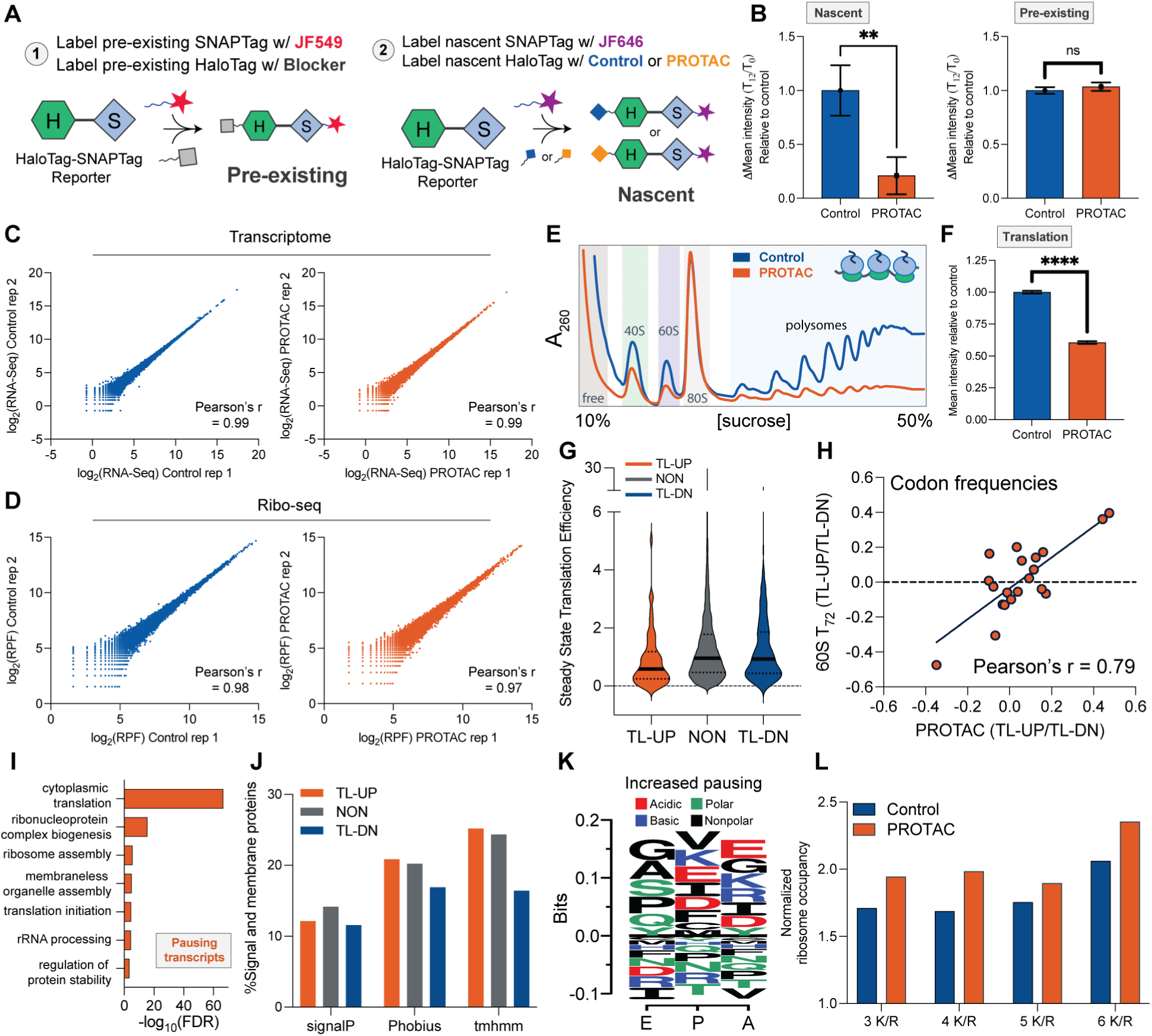
Validation and additional characterization of strategy to introduce cellular ribosome age imbalances. (A) Schematic of strategy used to quantify PROTAC-mediated degradation of nascent and pre-existing HaloTag fusions. (B) Quantification of nascent and pre-existing HaloTag degradation under Control and PROTAC treatment strategy described in (A) (n ≥ 178 cells, p ≤ 0.01, two-tailed Welch’s t test). (C,D) Correlations between replicates for transcriptome and Ribo-seq experiments under Control and PROTAC treatment. (E) Polysome analysis under Control and PROTAC treatment. (F) Quantification of translation activity under Control and PROTAC treatment by labeling of nascent polypeptides with OP-Puromycin (n ≥ 480 cells, p ≤ 0.0001, two-tailed Welch’s t test). (G) Steady state translation efficiencies for transcripts regulated by PROTAC treatment. (H) Correlation between codon frequency ratios for transcripts regulated by PROTAC treatment and molecularly aged 60S ribosomes. (I) Gene Ontology terms for pausing transcripts (adjusted *P* < 0.05). (J) Percent of signal and membrane proteins encoded by transcripts regulated by PROTAC treatment according to multiple algorithms. (K) Logo plot of amino acids being decoded at PROTAC-dependent pause sites. (L) Average ribosome occupancy at polybasic regions of different lengths under Control or PROTAC treatment.

**Figure S7.**
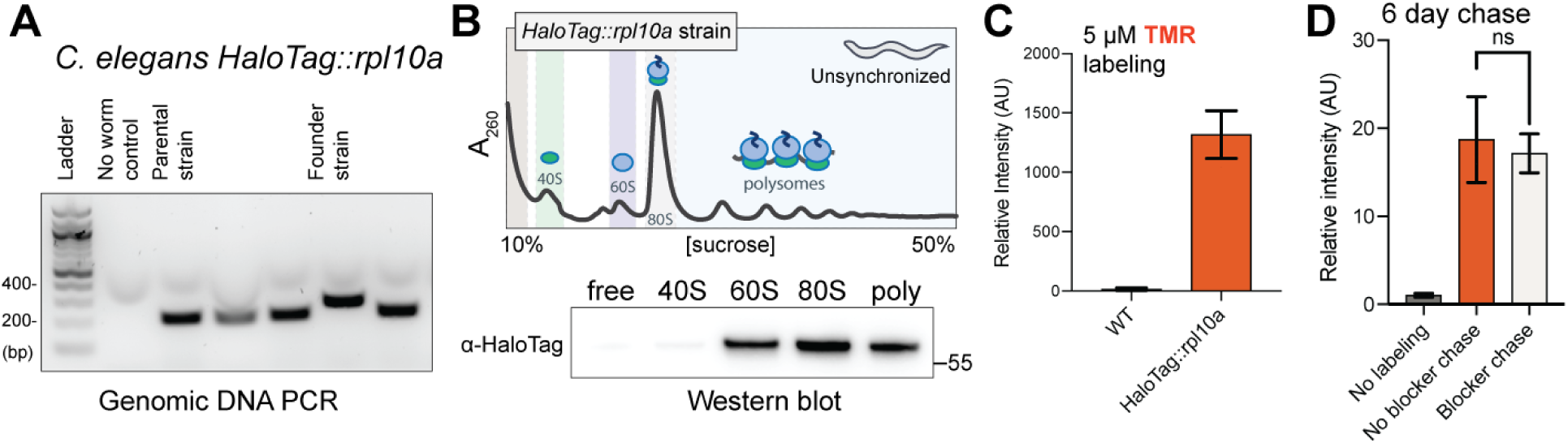
Characterization and pulse-chase labeling of *HaloTag::rpl10a C. elegans* strain. (A) Genomic DNA PCR of founder *HaloTag::rpl10a* strain used in this study. (B) Polysome analysis and western blot of distribution of HaloTag::RPL10A fusion over ribosome fractions in *C*. *elegans* strain. (C) Comparison of 5 μM HaloTag-TMR labeling in wild type vs. *HaloTag::rpl10a C. elegans* (n = 4). (D) Comparison of TMR ligand chase after 6 days in the presence or absence of HaloTag blocking ligand (n ≥ 6, p = ns, two-tailed Welch’s test).

## METHODS

### Cell lines and culture conditions

U2OS cells (ATCC, HTB-96) were cultured in DMEM (Gibco, 11995065) supplemented with 10% FBS (R&D Systems, S11150H), and 1% penicillin/streptomycin (Gibco, 15140122) in a humidified chamber at 37℃ with 5% CO_2_. For passaging, U2OS cells were dissociated with 0.05% Trypsin-EDTA (ThermoFisher, 25300054). HBEC3-KT cells (Normal Human Bronchial Epithelial Cells immortalized with CDK4 and hTERT, HBECs hereafter) were a gift from the Daniel Larson Lab (NCI). HBECs were grown in Keratinocyte-SFM medium (Gibco, 17005042) supplemented with 25 mg Bovine Pituitary Extract (BPE), 2.5 μg EGF, and 1% penicillin-streptomycin. Cells were dissociated with 0.05% Trypsin-EDTA and quenched with Trypsin Neutralizer Solution (ThermoFisher, R002100) before spinning down and passaging. Human patient fibroblasts (Coriell, Table S1) were cultured in EMEM (ATCC, 30-2003) supplemented with 20% FBS (Genesee Scientific, 25-550H) in a humidified chamber at 37℃ with 5% CO_2_. Cells were dissociated with Trypsin EDTA 1X Solution in HBSS (Fujifilm, 9341-500ML). For microscopy, cells were plated and imaged in 96-well glass bottom dishes (Cellvis, P96-1.5H-N). Cell growth assays were performed using the Incucyte system (Sartorius). For ribotoxicity tests, cells were treated with 300 nM anisomycin (SelleckChem, S7409) for 48 hours, then cell viability was measured with the MTT assay (Sigma Aldrich, 11465007001).

### Endogenous tagging of ribosomal subunits with HaloTag using CRISPR/Cas9 in U2OS cells

The termini of ribosomal proteins (RPL10A N-terminus, and RPS17 C-terminus) were endogenously tagged in U2OS cells via ribonucleoprotein (RNP) delivery of preassembled Cas9 protein (Alt-R™ S.p. HiFi Cas9 Nuclease V3, IDT), with guide RNAs, and single-stranded Alt-R™ HDR Donor Templates (IDT) with 200 bp homology sequences flanking HaloTag-Gly/Ser Linker sequences (Table S4). Guide RNAs were designed using CHOPCHOP v3.^92^ Guide RNA sequences used were TTCAGGACCAACTCACCTCA for RPL10A and ACTACAGAAAAAATTCAAAC for RPS17. Cells were nucleofected using the SE Cell Line 4D-Nucleofector X Kit S (Lonza, V4XC-1032) following the manufacturer’s protocol. To enhance the efficiency of homologous recombination, cells were incubated in media containing 30 μM Alt-R HDR enhancer (IDT) overnight, followed by changing to enhancer-free media. Cells were expanded, and monoclonal cell lines were isolated via single-cell FACS. Insertions were validated via western blotting against HaloTag, and junction PCR of the specific genomic locus (Figure S1A-E) using oligonucleotides synthesized by IDT, with subsequent sequence validation by Sanger sequencing (GENEWIZ).

### Endogenous tagging of SNRPB with HaloTag using CRISPR/Cas9 in HBECs

The N-terminus of SNRPB was endogenously tagged in wild-type HBEC3-KT cells with a CRISPR/Cas9 system (Figure S1F,G). The guide RNA, GAACCGCCACCATGGTAAGG, was designed with CRISPOR^93^ and cloned into a modified version of PX458 (Addgene, Plasmid #48138) with a Cas9-mCherry fusion (gift from the Daniel Larson Lab, NCI). The donor plasmid consisted of left and right homology arms each spanning ∼1kb, a HaloTag sequence, and a 3C protease cleavage site embedded in a Gly/Ser linker (Table S4). Cells were electroporated with 9 μg of donor template and 3 μg of Cas9-guide plasmid using the Neon electroporation system (Invitrogen, MPK5000). mCherry positive cells were sorted 48 hours after electroporation and expanded. Then, HaloTag-positive, monoclonal cell lines were isolated via single-cell FACS. Insertions were validated via imaging, western blotting (against both HaloTag and SNRPB), and PCR of the specific genomic locus using oligonucleotides synthesized by IDT.

### Plasmid construction and transfection

To construct DNA plasmids (Table S2) encoding emiRFP670-SEC61B, HaloTag, and NLS-HaloTag-SNAPTag, DNA fragments were amplified from original plasmids by PCR using Q5 High-Fidelity 2X Master Mix (NEB, M0494L) with oligonucleotides synthesized by IDT, then the In-Fusion HD Cloning kit (Takara, 638949) was used to insert the fragments into the FM5 vector^94^ under a Ubiquitin C promoter featuring a Gly/Ser-linker. The mTagBFP2-NPM1 construct was assembled as previously described.^95^ Previously described polybasic stalling reporters^96^ were modified by swapping GFP with mTagBFP2 by restriction digestion and insertion of synthetic sequences (GenScript) via the In-Fusion HD Cloning kit, resulting in mTagBFP2-P2A-K0-P2A-RFP, mTagBFP2-P2A-K12-P2A-RFP, and mTagBFP2-P2A-K22-P2A-RFP plasmids. Where applicable, plasmid DNA transfection of U2OS cells was carried out using FuGENE (Promega, PRE2311) according to the manufacturer’s instructions. All constructs were confirmed by Sanger sequencing (GENEWIZ).

### Lentiviral production and transduction

Lentivirus of respective plasmids were made using HEK293T cells seeded in a six-well plate at 70% confluence. The plasmids were co-transfected with helper plasmids VSVG and PSP using Lipofectamine 3000 (Invitrogen, L3000008) according to the manufacturer’s protocol. After 3 days, supernatant was harvested and filtered through syringe filters with a 0.45-µm pore size (VWR), then immediately used for transduction of U2OS cells plated at 30% confluency for 2 days. Cells were then expanded to make stable expression lines, and FACS-sorted using a tight window for each fluorescent reporter of interest.

### HaloTag ligand preparation and labeling of nascent and pre-existing proteins in human cells

HaloTag-TMR ligand was obtained from Promega (G8252). Non-fluorescent HaloTag blocking ligand was prepared by incubating 100 mM HaloTag succinimidyl ester (O4) ligand (Promega, P6751) with 500 mM Tris-HCl pH 8.0 (ThermoFisher, J22638.K2) for 60 min at 25℃ as previously described.^97^ For labeling of nascent proteins of different age distributions (‘Block-Label’), cells were incubated with 25 μM HaloTag blocking ligand in complete media at 37℃ for 30 minutes to achieve saturation of HaloTag binding sites. Cells were then washed once with DPBS, then three times with ligand-free complete media. Following this, cells were labeled with 250 nM HaloTag-TMR ligand continuously for the desired time intervals at 37℃. For labeling of pre-existing proteins of different age distributions (‘Label-Block’), cells were first incubated with 250 nM HaloTag-TMR ligand in complete media at 37℃ for 30 minutes to achieve saturation. Cells were then washed once briefly with DPBS, then three times with ligand-free complete media at 37℃ for 5 minutes each. Following this, cells were incubated with 25 μM HaloTag blocking ligand in complete media continuously at 37℃ for the desired time intervals. SNAPTag-JF549 and -JF646 ligands were obtained from HHMI Janelia. Labeling optimization is described in Figure S1I,J.

### Confocal microscopy of living cells

Images were acquired on a Nikon CSU-W1 SoRa spinning-disc confocal microscope equipped with a Yokogawa SoRa pixel reassignment-based super-resolution device and dual Hamamatsu Fusion BT sCMOS cameras. All acquisitions were performed using a Nikon CFI Plan Apo Lambda D 60X oil (MRD71670) lens, and 405, 488, 561 and 640 nm lasers were used where required. Denoise.ai (Nikon software) was used for images shown and analysed in this study. During imaging, cells were maintained in a humidified chamber at 37℃ with 5% CO2.

### Preparation of anti-TMR antibody-coated magnetic beads

Anti-TMR beads were prepared by covalently coupling 15 μg of TRITC polyclonal antibody (ThermoFisher, A-6397) for every 1 mg of Dynabeads M-270 Epoxy beads (ThermoFisher, 14301) according to the manufacturer’s instructions and stored at 4℃ in SB buffer included in the kit until use.

### Immunopurification of nuclear pre-60S intermediates

Pre-60S ribosomal subunit subpopulations of interest were labeled with the HaloTag-TMR ligand as described above. For each immunopurification (IP) experiment, two 15 cm plates of HaloTag-RPL10A or HaloTag-only U2OS cells (Figure S1H) at ∼70% confluency were used. All experiments were done in biological triplicates. Cell lysis conditions were adapted from previously described work.^44^ Cells were washed twice with ice-cold DPBS supplemented with calcium and magnesium (ThermoFisher, 14040133), then lysed in Buffer NA composed of 25 mM HEPES pH 7.6 (ThermoFisher, J61047.AE), 65 mM NaCl (ThermoFisher, AM9760G), 65 mM KCl (ThermoFisher, AM9640G), 3 mM MgCl2 (ThermoFisher, AM9530G), 1 mM EDTA (ThermoFisher, 15575020), 10% Glycerol (ThermoFisher, 327255000), 0.05% Triton X-100 (Sigma Aldrich, T8787), 1 mM DTT (ThermoFisher, 707265ML), 1X cOmplete EDTA-free protease inhibitor (Sigma Aldrich, 4693132001) in RNase/DNase-free water (Fisher Scientific, 10-977-023), and dounced 7 times with an RNase-free pestle (Fisher Scientific, 12-141-364) on ice to lyse. The lysate was centrifuged at 20,000g, for 5 minutes at 4℃. The resulting pellet was washed twice with Buffer NA, centrifuging at 20,000g, for 5 minutes at 4℃ each time. The resulting pellet was then incubated in Buffer NB, composed of 25 mM HEPES pH 7.6, 10 mM KCl, 10 mM MgCl2, 1 mM CaCl2 (Sigma Aldrich, 21115), 1 mM EDTA, 100 mM Arginine (Sigma Aldrich, A5006), 25 mM ATP (ThermoFisher, J61125), 1 mM Spermidine (Sigma Aldrich, S0266), 5% glycerol, 0.1% Triton X-100, 1 mM DTT, 1X cOmplete EDTA-free protease inhibitor, 25U/mL TurboDNase (ThermoFisher, AM2239), 1:40 Murine RNase Inhibitor (NEB, M0314L) in in RNase/DNase-free water, on a nutator for 30 minutes at 4℃ to lyse. The lysate was centrifuged at 20,000g for 20 minutes at 4℃ to remove the insoluble fraction. Protein concentrations from the resulting lysates were measured using the Qubit Protein Assay (Thermo Fisher Scientific, Q33211) with a Qubit fluorometer, and concentrations were normalized. For each IP, 3 mg of anti-TMR beads resuspended in Buffer NB were added and incubated for 4 hours on rotation at 4℃. After this, beads were washed twice on ice with Buffer NC composed of 25 mM HEPES pH 7.6, 75 mM NaCl, 75 mM KCl, 5 mM MgCl2, 0.5 mM EDTA, 100 mM Arginine, 2.5% glycerol, 0.1% Triton X-100, 1 mM DTT, 1X cOmplete EDTA-free protease inhibitor in RNase/DNase-free water. Then, beads were washed once with Buffer ND composed of 25 mM HEPES pH 7.6, 75 mM NaCl, 75 mM KCl, 5 mM MgCl2, 0.5 mM EDTA, 100 mM Arginine, 2.5% Glycerol, 0.05% C12E8 (Anatrace, O330), 1X cOmplete EDTA-free protease inhibitor in RNase/DNase-free water. To elute proteins off the anti-TMR beads, samples were incubated at 85℃ for 10 minutes with shaking at 1000 rpm in 1X TES buffer composed of 300 mM Tris-HCl pH 8.0, 1% SDS (ThermoFisher, AM9820), 0.5 mM EDTA in water.

### Immunopurification of cytoplasmic ribosome subpopulations

Ribosome subpopulations of interest were labeled with the HaloTag-TMR ligand as described above. For each immunopurification (IP) experiment, two 15 cm plates of either HaloTag-RPL10A, RPS17-HaloTag, or HaloTag-only control U2OS cells at ∼70% confluency were used. All experiments were done in biological triplicates. Cell lysis conditions were adapted from previously described work.^98^ To stabilize polysomes, cells were treated with 100 μg/mL cycloheximide (Fisher Scientific, AAJ66004XF) for 2 minutes at 37℃ prior to lysis. Cells were washed twice with ice-cold DPBS supplemented with calcium and magnesium, and 100 μg/mL cycloheximide, then lysed in Buffer A, composed of 25 mM HEPES pH 7.3 (ThermoFisher, J16924.AE), 150 mM NaCl, 15 mM MgCl2, 8% glycerol, 1% Triton X-100, 0.5% sodium deoxycholate (Sigma Aldrich, 30970), 100 μg/mL cycloheximide, 1 mM DTT, 25 U/mL TurboDNase, 1:40 Murine RNase Inhibitor, 1X cOmplete EDTA-free protease inhibitor, in RNase/DNase-free water, and dounced 7 times with an RNase-free pestle on ice to lyse. Lysates were depleted of nuclei via successive centrifugations at 4℃ for 5 minutes each: twice at 800g, once at 8,000g and once at 20,000g, keeping the supernatant from each centrifugation. Protein concentrations from the resulting lysates were measured using the Qubit Protein Assay (Thermo Fisher Scientific, Q33211) with a Qubit fluorometer, and concentrations were normalized. For each IP, 3 mg of anti-TMR beads resuspended in Buffer A were added and incubated for 3 hours on rotation at 4℃. After this, beads were washed three times with Buffer B composed of 25 mM HEPES pH 7.3, 150 mM NaCl, 15 mM MgCl2, 1 mM DTT, 1% Triton X-100, 0.5% sodium deoxycholate, 100 μg/mL cycloheximide, 1X cOmplete EDTA-free protease inhibitor, in RNase/DNase-free water, and three times with Buffer C composed of 25 mM HEPES pH 7.3, 300 mM NaCl, 15 mM MgCl2, 1 mM DTT, 1% Triton X-100, 0.5% sodium deoxycholate, 100 μg/mL cycloheximide, 1X cOmplete EDTA-free protease inhibitor, in RNase/DNase-free water. To elute proteins off the anti-TMR beads, samples were incubated at 85℃ for 10 minutes with shaking at 1000 rpm in 1X TES buffer.

### Immunopurification of spliceosomes

Spliceosomes were labeled with the HaloTag-TMR ligand as described above. For each immunopurification (IP) experiment, three 15 cm plates of either HaloTag-SNRPB, or HaloTag-only control HBEC cells at ∼70% confluency were used. All experiments were done in biological triplicates. Cell lysis conditions were adapted from previously described work.^99^ Cells were washed twice with ice-cold DPBS supplemented with calcium and magnesium, then harvested in PBS by scraping with an RNase-free cell scraper (CellTreat, 229310). Cells were pelleted by centrifugation at 300g for 5 minutes at 4℃, and washed twice with cytoplasmic depletion buffer composed of 20 mM HEPES pH 8.0 (ThermoFisher, J63002.AK), 65 mM NaCl, 65 mM KCl, 3 mM MgCl2, 1 mM EDTA, 10% Glycerol, 0.5% Triton X-100, 1 mM DTT, 50 U/mL SUPERase In RNase Inhibitor (ThermoFisher, AM2696), 1X cOmplete EDTA-free protease inhibitor, in RNase/DNase-free water, pelleting nuclei by centrifugation at 20,000g for 5 minutes at 4℃ between each wash. The resulting nuclei were lysed for 10 minutes on ice in EJC lysis buffer composed of 20 mM Tris-HCl pH 7.5 (ThermoFisher, 15567027), 15 mM NaCl, 10 mM EDTA, 0.5% NP-40 (Sigma Aldrich, 74385), 0.1% Triton X-100, 1.0 mM DTT, 1X cOmplete EDTA-free protease inhibitor tablet, 100 U/mL SUPERase In RNase inhibitor in RNase/DNase-free water. Nuclear pellets were disrupted by douncing with an RNase-free pestle, and the suspension was sonicated at 15% amplitude using a micro-tip sonicator (Branson) for a total of 16 seconds (2 s bursts with 10 s intervals). The NaCl concentration was adjusted to 150 mM, then the lysate was cleared via centrifugation at 10,000g for 10 minutes at 4℃. RNA concentrations from the resulting lysates were measured using the Qubit RNA Broad Range (BR) assay (Thermo Fisher Scientific, Q10211) with a Qubit fluorometer, and RNA concentrations were normalized. For each IP, 3 mg of anti-TMR beads resuspended in Buffer EJC adjusted to 150 mM NaCl were added and incubated for 3 hours on rotation at 4℃. After this, beads were washed twice with isotonic wash buffer composed of 20 mM Tris-HCl pH 7.5, 150 mM NaCl, 0.1% NP-40, 1.0 mM DTT, 1X cOmplete EDTA-free protease inhibitor tablet in RNase/DNase-free water. Then, beads were washed four times with Buffer WB225 composed of 20 mM Tris-HCl pH7.5, 225 mM NaCl, 0.1% NP-40, 1.0 mM DTT, 1X cOmplete EDTA-free protease inhibitor tablet, in RNase/DNase-free water. To elute off the anti-TMR beads, samples were incubated at 85℃ for 10 minutes in 1X TES buffer.

### Sample preparation for mass spectrometry analysis

Eluates were first adjusted to 3% SDS (w/v), then reduced and alkylated via the addition of 25 mM TCEP and 50 mM chloroacetamide at 70℃ for 20 min. Samples were then acidified by addition of phosphoric acid to a final concentration of 1.2% (w/v). Samples were prepared for trypsin digestion ahead of LC/MS analysis with S-trap micro spin columns (Protifi, C002-MICRO) following manufacturer recommendations. On-column trypsin digestion was performed at 37℃ for 16 hours. Following trypsinization, samples were sequentially eluted in 25 mM TEAB, 0.1% formic acid, and 50% acetonitrile/0.1% formic acid, then dried via speed-vac. Samples analyzed on a Q-Exactive HF instrument (ThermoFisher scientific), including nuclear pre-60S, cytoplasmic 60S and cytoplasmic 40S purifications, were resuspended in 1% acetonitrile/0.1% formic acid to a final concentration of 0.75 μg/μL, while samples analyzed on a timsTOF HT (Bruker), including spliceosome and nascent polypeptide purifications, were resuspended in 4% acetonitrile/0.1% formic acid/0.15% n-dodecyl-ß-D-maltoside (DDM) to a final concentration of 100 ng/μL.

### Mass spectrometry data acquisition

For samples analyzed on a Q-Exactive HF mass spectrometer equipped with a Nanospray Flex Ion Source (ThermoFisher Scientific), 1.5 µg of peptides were resolved for nLC-MS analysis with a Dionex Ultimate 3000 nRSLC (ThermoFisher Scientific) equipped with an EASY-Spray HPLC column (ThermoFisher ES803, 50 cm x 75 µm ID, PepMap RSLC C18, 2 µm pore size). Peptides were resolved with mobile phases 0.1% formic acid (A) to 0.1% formic acid in 97% acetonitrile (B) over a two-step 80-minute gradient (0-25% over 60 minutes, 25-50% over 20 minutes) using with a flow rate of 0.25 µL/min. MS and MS/MS spectra were automatically obtained in a full MS/data-independent MS2 method. The MS1 method operated with a full scan range of 395-1005 *m/z* with a resolution of 60,000, a target AGC of 3e6 with a maximum injection time (MIT) of 55 ms, and spectra were recorded in centroid. For data-independent MS2 scans, an isolation window of 24.0 m/z was used, with a resolution of 30,000, an AGC target of 3e6, a loop count of 25, maximum injection time of 55 ms, and collision energy of 28.

For samples analyzed on a Bruker timsTOF HT equipped with a CaptiveSpray 2 ion source, 100 ng of peptides were resolved for nLC-MS analysis with a nanoElute2 autosampler (Bruker) coupled with a PepSep Ultra C18 column (15 cm x 75 µm ID, C18, 1.5 µm pore size). Peptides were resolved with mobile phases 0.1% formic acid (A) to 0.1% formic acid in acetonitrile (B) over a linear 60-minute gradient (0-35%) using a flow rate of 0.3 µL/min. Intact peptide and fragment ion measurements were acquired using a data independent acquisition-parallel accumulation serial fragmentation (dia-pasef) method.^100^ dia-pasef isolation windows were optimized using pydiaid,^101^ followed by manual adjustment of ion mobility boundaries. A scan range of 100 m/z–1,700 m/z with a 0.60 Vs/cm^2^–1.57 Vs/cm^2^ mobility (1 /_K0_) range was used with a 100 ms ramp time and a collision energy of 10 eV.

### DIA data analysis

DIA-NN^102^ was used to search raw MS data with the following settings. First, a FASTA file containing the human reference proteome (UP000005640) was amended to also include rabbit IGG (P01870, P01879), haloalkane dehalogenase (P0A3G3), and streptavidin (P22629). From these FASTA files, an *in silico* spectral library was created. Analysis allowed for one missed cleavage, with N-terminal M excision and cysteine carbamidomethylation as modifications. Peptides between 7 and 30 amino acids in length with a precursor charge of 1–4 (QE-HF) or 2-4 (timsTOF HT) and with precursor and fragment m/z values within the range of 300–1,800 and 200–1,800, respectively, were quantified. MS1 and MS2 mass accuracy were both set to 15.0. We used a high-precision quantification strategy with double-pass mode for the neural network classifier. For maximum parsimony, match between runs and cross-run normalization were turned off.

### Proteomics data normalization

FragPipe Analyst^103^ was used to identify bona fide interaction partners relative to control samples. HaloTag-based immunopurification proteomics datasets were normalized within each sample to bait (HaloTag) abundance. For HaloTag immunopurifications, steady-state (total labeling) samples were compared against HaloTag-only controls. For nascent polypeptide purifications, O-propargyl-puromycin-labeled cells were compared with unlabeled controls. Differential expression analysis was performed using an adjusted p-value cutoff of 0.05 and a log_2_ fold-change threshold of 1. Missing values were imputed using a Perseus-type imputation strategy. Multiple hypothesis testing correction was applied using the Benjamini-Hochberg false discovery rate (FDR) method. The minimum required percentage of non-missing values was set to 0% globally and 66% within at least one condition. For HaloTag immunopurification experiments, all steady-state and pulse-chase timepoint samples were internally normalized by a HaloTag peptide that was consistently detected across all samples. Data in Figure 1F were fit by nonlinear least-squares regression using a four-parameter logistic (4PL) sigmoidal model in GraphPad Prism, with concentration as the independent variable and response as the dependent variable.

### Total RNA isolation and electrophoresis

Cells were collected in 1X Buffer RLT (QIAGEN, 79216) and isolated using the QIAGEN RNeasy Mini Kit (74104) according to the manufacturer’s protocol. The samples were treated with Turbo DNase (Thermo Fisher Scientific, AM2238) at 37℃ for 1 hr. DNase-digested RNA samples were subsequently purified using the Zymo RNA Clean and Concentrator-25 kit (R1017). RNA integrity after isolation and 28S to 18S rRNA ratio quantification where required were assayed using the Agilent RNA High Sensitivity Assay (Agilent, 5067-5579) on the 4150 TapeStation system (Agilent Technologies) according to the manufacturer’s protocol.

### Western blotting

Cells were harvested by rinsing with ice-cold 1X DPBS and lysing in 1X RIPA buffer for 30 minutes on ice. Lysates were centrifuged at maximum speed, and supernatants were collected for subsequent assays. The Pierce Detergent Compatible Bradford Assay Kit (ThermoFisher, 23246) was used to determine protein concentration and normalize each sample. Lysates were denatured in a NuPAGE LDS Sample Buffer (ThermoFisher, NP0007), supplemented with NuPAGE™ Sample Reducing Agent (ThermoFisher, NP0004) at 95℃ for 5 minutes. Normalized protein samples were loaded onto NuPAGE 4-12% Bis-Tris Protein Gels (ThermoFisher, NP0322) and run for 20 minutes at 80V then for 75 minutes at 120V in NuPAGE MES SDS Running Buffer (ThermoFisher, NP0002). Proteins were then transferred to 0.2 μm nitrocellulose membranes (BioRad, 1704158) on the Trans-Blot Turbo System (BioRad, 17001917). Membranes were blocked in blocking buffer containing 1X TBST (BioRad, #1706435) buffer and 5% bovine serum albumin (R&D systems, 5217) for 30 minutes, then incubated with primary antibodies overnight at 4℃ in blocking buffer. All primary antibodies were used at 1:1000 dilution. The primary antibodies used were: anti-HaloTag (Promega, G9211), anti-TRITC, anti-RPL10A (ThermoFisher, MA5-44710), anti-RPS17 (ThermoFisher, PA5-95760), anti-SNRPB (ThermoFisher, MA513449), anti-RPS18 (Novus Biologicals NBP2-93632), anti-RPL5 (Abcam, ab86863), anti-RPL9 (Novus Biologicals NBP1-82853), anti-EIF4A1 (Cell Signaling Technology, 2490T) and anti-EEF1A (Cell Signaling Technology, 2551S). After primary antibody incubation, blots were washed with 1X TBST buffer three times (5 minutes per wash), then stained with secondary antibodies at 1:5,000 dilution for 1 hr at room temperature. The secondary antibodies used were: Peroxidase AffiniPure Goat Anti-Rabbit IgG (H+L) (Jackson Immunology, 111-035-045), and Peroxidase AffiniPure™ Goat Anti-Mouse IgG (H+L) (Jackson Immunology, 115-035-062). The blots were washed three times with 1X TBST buffer for 5 minutes each time before developing with Clarity Western ECL Blotting Substrates (BioRad, 1705061) and imaging on the ChemiDoc Imaging System (BioRad, 12003153).

### Polysome analysis in human cells

Cells were seeded in 15 cm plates at 70% confluence at the day of experiment. Prior to harvest, cells were treated with 100 μg/mL cycloheximide in complete media for 2 minutes at 37℃ to stabilize polysomes. Cells were washed twice with ice-cold DPBS supplemented with calcium and magnesium, then lysed in Buffer A. Lysates were depleted of nuclei via successive centrifugations at 4℃ for 5 minutes each: twice at 800g, once at 8,000g and once at 20,000g, keeping the supernatant from each centrifugation. RNA concentrations from the resulting lysates were measured using the Qubit RNA Broad Range (BR) assay with a Qubit fluorometer, and RNA concentrations were normalized. Normalized lysates were layered onto a 10%-50% sucrose gradient prepared from 1X polysome buffer containing 25 mM HEPES pH 7.3, 150 mM NaCl, 15 mM MgCl2, 1 mM DTT, 100 μg/mL cycloheximide, 1X cOmplete EDTA-free protease inhibitor in RNase/DNase-free water. The samples were then centrifuged in a SW41Ti rotor (Beckman Coulter) for 2.5 hr at 40,000 rpm at 4℃. Following centrifugation, the samples were continuously recorded and fractionated into individual tubes on a Biocomp Gradient Fractionator, equipped with a Triax flow cell and a Gilson fraction collector according to the manufacturer’s instructions. For analysis of proteins from polysome fractions, proteins were precipitated using the ProteoExtract Protein Precipitation Kit (Fisher Scientific, 53-918-01KIT) according to the manufacturer’s instructions. Protein concentrations were determined with the Pierce Detergent Compatible Bradford Assay Kit and fractions normalized, then characterized by western blotting.

### Generation of *C. elegans* strains

CRISPR knock-in of HaloTag as an N-terminal fusion at the endogenous *rpl-10a* locus in wild-type background was performed by SunyBiotech. Due to homozygous sterility, a subsequent balancer cross generated strain PHX9377, rpl-10a(syb9377)/hT2 [bli-4(e937) let-?(q782) qIs48] (I;III). A further CRISPR edit was performed as previously described^104^ to lengthen the glycine-serine linker from four to 22 amino acids (Table S4), confirmed by PCR genotyping and Sanger sequencing. This *HaloTag::rpl-10a* strain propagates as a mixture of homozygotes and balanced heterozygotes at ∼1:4 ratio.

### *C. elegans* synchronization and culture

Synchronized *HaloTag::rpl-10a* populations were generated by harvesting gravid hermaphrodites from 40 6 cm NGM plates via S-basal wash. Animals were subjected to alkaline hypochlorite lysis (1% NaOCl, 0.5M NaOH) to isolate embryos, which were washed three times in S-basal and transferred to NGM plates seeded with *E. coli* OP50 and grown at 20℃. Upon reaching the L4 stage, 60,000 animals were transferred to 10 cm seeded NGM plates supplemented with 50 μg/mL 5-fluoro-2’-deoxyuridine (FUdR ; MedChemExpress HY-B0097) to inhibit progeny production. Populations were maintained *ad libitum* with concentrated OP50 and monitored daily to ensure well-fed conditions through Day 10.

### Ribosome pulse-chase labeling in *C. elegans*

For HaloTag-TMR labeling of ribosomes, 40,000 worms were labeled from Day 1 to Day 2 of adulthood in 50 mL conical tubes, with 10 mL total culture volume at a density of ∼1 worm/μL. Labeling was performed in S-basal containing 10x concentrated OP50, 50 μg/mL FUdR, and 5 μM HaloTag TMR ligand on a tube rotator for 24 hours. Following incubation, residual ligand was removed via six serial washes in S-basal with 50 μg/mL FUdR. Animals were grown on FUdR-supplemented seeded NGM plates until Day 9, then transferred to liquid culture. On Day 10, worms were harvested and washed in M9 buffer, and the resulting pellets were flash-frozen in liquid nitrogen and stored at -80℃.

### Polysome analysis in *C. elegans*

For polysome analysis in *C.* elegans, 2,500 worms flash-frozen in liquid nitrogen and stored at -80℃ were thawed in 350 μL worm polysome buffer (25 mM Tris-HCl pH 7.5, 140 mM KCl, 1.5 mM MgCl2, 200 μg/ml cycloheximide, 1% Triton X-100, 0.1% sodium deoxycholate, 1 mM DTT, 1x cOmplete EDTA-free protease inhibitor, 5U/mL TurboDNase) and immediately homogenized using RNase-free pestles, using 25 strokes. For analysis excluding nuclease treatment, 200 U SUPERase•In RNase Inhibitor was added to the worm polysome buffer. The homogenized worms were incubated for 30 minutes at 4℃ on a nutator to allow lysis to occur, then centrifuged at 16,000g for 10 minutes at 4℃ to remove debris. RNA concentrations were measured using the Qubit RNA Broad Range (BR) assay with a Qubit fluorometer and normalized. For nuclease treatment, 150 U RNase I (Ambion, AM2294) per 200 μg of RNA was added, and incubated for 30 minutes on rotation at room temperature, followed by quenching by adding 200 U SUPERase•In RNase Inhibitor. Lysates were then layered and analyzed onto 10%-50% sucrose gradients as described above. For analysis of proteins from polysome fractions, proteins were precipitated using the ProteoExtract Protein Precipitation Kit (Fisher Scientific, 53-918-01KIT) according to the manufacturer’s instructions. Protein concentrations were determined with the Pierce Detergent Compatible Bradford Assay Kit and fractions normalized, then characterized by western blotting. Steady state ribosomes were probed via the anti-HaloTag antibody (1:1000), and aged ribosomes were probed via the anti-TMR (TRITC) antibody (1:1000).

### RNase I treatment for human cell samples

Buffer A excluding RNase inhibitors was used to lyse cells. RNA concentrations from clarified cell lysates were quantified using the Qubit RNA Broad Range (BR) assay with a Qubit fluorometer and normalized. 150 U RNase I were added for every for 30 μg of RNA in a 400 μL volume and incubated for 30 minutes at room temperature with end-to-end rotation. The RNase reaction was quenched by adding 10 μL of 20U/μL SUPERase In RNase Inhibitor. Digested lysates were layered on top of sucrose gradients as described above.

### Purification of RNA from age-specific ribosomes from human cells

HaloTag-RPL10A or RPS17-HaloTag U2OS cells were incubated with 250 nM HaloTag-TMR ligand in complete media at 37℃ for 30 minutes to label all ribosomes to saturation. Cells were then washed once briefly with DPBS, then three times with ligand-free complete media at 37℃ for 5 minutes each. Following this, cells were incubated with 25 μM HaloTag blocking ligand in complete media continuously at 37℃ for the following time intervals: 0 (‘steady-state’), 24 and 72 hr. Cells were replenished with fresh media daily. For each experiment, six 15 cm plates were used at a final confluency of ∼70% at the time of harvest. All experiments were done in biological duplicates. For experiments on fractionated lysates, cells were lysed in Buffer A excluding RNase inhibitors and RNase I treatment was performed as described above. Lysates were then fractionated on 10%-50% sucrose gradients, and the monosome (80S) or disome fractions were kept for further processing depending on the experiment. For experiments without fractionation, cells were lysed in Buffer A including 1:40 Murine RNase Inhibitor and immediately followed by immunopurification. The fractions were diluted threefold with Buffer B, after which 4 mg of anti-TMR beads resuspended in Buffer B were added, then incubated for 3 hours at 4℃ with rotation. The beads were then washed four times with Buffer B, then four times with Buffer C. RNAs were eluted off the anti-TMR beads by incubating in NLS buffer composed of 2% N-lauryl sarcosine (Milipore Sigma, L7414), 10 mM EDTA and 1X PBS, supplemented with 8 U Proteinase K (NEB, P8107S) and incubated at 50℃ for 1 hr with shaking at 1000 rpm. RNAs were then purified using the Oligo Clean & Concentrator kit (Zymo, D4061).

### Analysis of cell cycle distribution

Cells were subjected to either the T0 or T72 labeling procedure. In the T72 protocol, cells were incubated with 25 μM HaloTag blocking ligand continuously throughout the 72-hour chase period with daily media replenishment. Cells were then incubated with 10 μM EdU (Thermo Fisher Scientific, C10340) in complete culture medium for 1 hour at 37℃ to label S-phase cells undergoing DNA replication. Following EdU incorporation, cells were trypsinized and pelleted at 500g for 5 minutes, washed once with PBS, and pelleted again. Cells were fixed in 4% PFA for 12 minutes at room temperature, pelleted, and washed once with PBS. Samples were permeabilized with 0.5% Triton X-100 in PBS (PBST) for 15 minutes at room temperature. EdU was detected by clicking AlexaFluor-647 according to the manufacturer’s instructions (Thermo Fisher Scientific, C10340). Cells were washed with PBS and pelleted. Total DNA was stained with DAPI (Thermo Fisher Scientific, D1306) at 2.5 μg/mL for 10 minutes at room temperature. Single-color control samples were prepared in parallel and used for compensation and gating. Flow cytometry was performed on a FACSymphony A3 (BD Biosciences), and data was analyzed using the FlowJo software.

### Labeling of newly synthesized ribosomal subunits with HaloPROTAC3

HaloTag-RPL10A U2OS cells were labeled to saturation with 250 nM HaloTag-TMR ligand for 30 min at 37℃ in complete medium. Cells were washed once with DPBS followed by three washes with ligand-free complete medium for 5 minutes each, and then incubated continuously at 37℃ in complete medium containing either 1.0 μM HaloPROTAC3 (PROTAC; Promega GA3110) or 1.0 μM ent-HaloPROTAC3 (Control; Promega GA4110) for the indicated time intervals. For Ribo-seq experiments, cells were treated with either PROTAC or control ligand for 72 hr, with daily replenishment of complete medium. For validation experiments using the NLS-HaloTag-SNAPTag reporter (Figure S6A,B), U2OS cells expressing the reporter were first labeled to saturation with both 25 μM HaloTag blocking ligand and SNAPTag JF549 ligand to block pre-existing binding sites. Then, cells were labeled continuously with either 1.0 μM HaloPROTAC3 or 1.0 μM ent-HaloPROTAC3, plus SNAPTag-JF646 ligand to track newly made proteins, then analyzed by microscopy after 12 hours of labeling.

### Generation and validation of Ψ18S-210 overexpression and control cells

SNORA10 was cloned into intron 1 of RPL23A, replacing the endogenous snoRNA. Because SNORA10 directs two distinct pseudouridylation events, Ψ28S-4491 and Ψ18S-210, and our goal was to selectively overexpress Ψ18S-210, we specifically inactivated the Ψ28S-4491 modification activity. To achieve this, the 28S rRNA guide sequence within SNORA10 was mutated to its reverse complement, generating the SNORA10-28Smut variant. The modified snoRNA sequence inserted into a two-exon EGFP sequence as previously described^105^ was synthesized (GenScript) and subsequently cloned into the pHR vector under the control of the SFFV promoter using the In-Fusion HD Cloning kit. An otherwise identical plasmid lacking the snoRNA insert served as a control. Lentiviral particles were used to stably transduce U2OS cells, which were then FACS-sorted to obtain populations with comparable EGFP expression levels. Expression of SNORA10-28Smut was confirmed by RT-qPCR (Figure S5E). Exclusive Ψ18S-210 modification activity and relative overexpression levels were assessed using BIHIND-qPCR^106^ according to the original protocol (Table S3, Figure S5F). Wild type SNORA10 expression in human patient fibroblasts (Figure S5L) was measured by RT-qPCR.

### Labeling and purification of nascent polypeptides

Nascent polypeptides were labeled by treating cells with 2 μM O-propargyl-puromycin (OPP, MedChemExpress HY-15680). For imaging-based translation measurements, cells were treated with OPP for 60 minutes at 37℃ in complete medium. Cells were then fixed with 4% paraformaldehyde (PFA, Electron Microscopy Science 15710) for 12 minutes at room temperature. Next, OPP was labeled with the Click-iT™ Plus Alexa Fluor™ 647 Picolyl Azide Toolkit (Thermo Fisher, C10643) according to the manufacturer’s instructions. Purification of nascent polypeptides labeled with OPP was adapted from previous work.^107^ Cells seeded at 70% confluence were labeled with OPP for 60 minutes at 37℃ in complete medium. After this, cells were harvested and lysed in 1X RIPA buffer supplemented with 1X cOmplete EDTA-free protease inhibitor. Lysates were cleared by centrifugation at 20,000g for 20 minutes at 4℃, and protein concentrations were measured and normalized using the Qubit Protein Assay with a Qubit fluorometer. Then, lysates were incubated for 2 hours at 4℃ on rotation with 50 μL Dynabeads MyOne Streptavidin C1 (ThermoFisher, 65002) to deplete endogenously biotinylated proteins prior to click chemistry. To biotinylate OPP-labeled nascent polypeptides, the resulting lysates were incubated for 1.5 hours at room temperature with 1 mM biotin picolyl azide (Click Chemistry Tools, 1167-25), 12 mM sodium ascorbate (Millipore Sigma, PHR1279), 10 mM THPTA (Millipore Sigma, 762342) and 2 mM CuSO4 (Millipore Sigma C1297). The reaction was quenched by precipitating proteins using the ProteoExtract Protein Precipitation Kit, incubating overnight at -20℃. Purified protein pellets were resuspended in 1X RIPA buffer supplemented with 1X cOmplete EDTA-free protease inhibitor. Streptavidin bead capture followed by on-bead trypsin digestion was then performed as described previously.^108^

### RNA-seq library preparation

RNA-Seq libraries were prepared as previously described.^109^ The final libraries were sequenced using 50 bp paired-end reads on an Illumina NovaSeq SP sequencer.

### Ribo-seq library preparation

Ribo-seq^110,111^ library preparation was performed as previously described^112^ with modifications. RNA obtained from sucrose gradient fractionations followed by molecular age-selective immunopurifications was denatured at 80℃ for 90 s and resolved on 15% TBE-urea polyacrylamide gels (ThermoFisher, EC68852BOX) in RNase-free 1X TBE buffer at 200 V for 65 min. To isolate ribosome footprints, gel fragments corresponding to 25-30 nt were excised. Footprints were eluted from the gel slices by freeze-thaw extraction (dry ice freeze, then thaw at room temperature overnight) in 400 μL RNA extraction buffer (300 mM sodium acetate, pH 5.2 (Sigma Aldrich, 567422); 1 mM EDTA; 0.25% (v/v) SDS). The eluate was collected, mixed with 1.5 μL GlycoBlue (ThermoFisher, AM9515) and 500 μL isopropanol, then frozen at -20℃ overnight. The RNA was pelleted by centrifugation at 20,000g for 30 min at 4 °C. Pellets were air-dried and resuspended in 8 μL nuclease-free water. 3.5 μL of eluted ribosome footprints were dephosphorylated in a 5 μL reaction containing 0.5 μL 10X T4 PNK buffer (NEB), 0.5 μL 10 U/μL T4 PNK enzyme (NEB, M0201L), and 0.5 μL SUPERase In RNase Inhibitor, incubated for 60 min at 37℃. Then, a 5 μL ligation reaction mixture composed of 3.5 μL 50% (w/v) PEG-8000 (NEB), 0.5 μL 10X T4 RNA ligase buffer and 0.5 μL 20 μM 5’ pre-adenylated linkers (Table S3) and 0.5 μL 200 U/μLT4 Rnl2(tr) K227Q (NEB, M0351S) was directly added, mixed, and incubated for 3 hours at 22℃, shaking at 1,600 rpm with 1 min on and 5 min off cycles. To specifically deplete unligated linkers from the reaction before reverse transcription, 1 μL 50 U/μL yeast 5’-deadenylase (NEB, M0331S) and 1 μL 10 U/μL RecJ exonuclease (Biosearch technologies, RJ411250) were added directly to the ligation reaction and incubated for 90 minutes at 30℃, with continuous shaking at 1,100 rpm, then purified using the Oligo Clean & Concentrator kit (Zymo) following the manufacturer’s instructions and eluted in 11 μL nuclease-free water. Ribosomal RNA depletion was skipped. For reverse transcription (RT), 10 μL RNA sample was mixed with 2 μL reverse transcription primer (NI-802, 1.25 μM) and denatured for 5 min. at 65℃. Then, 8.5 μL of RT master mix composed of 4 μL 5X First Strand (FS) Buffer, 1 μL 0.1 M DTT, 2 μL 10 mM dNTP mix, 0.5 μL RNase inhibitor, and 1 μL SuperScript III (ThermoFisher, 18080085) was added on ice, then incubated at 55℃ for 30 minutes, then held at 4℃. To clean-up the reaction, 4 μL Exo-SAP IT (ThermoFisher, 75001.1.ML) was added and incubated for 30 min at 37℃. Then, 1 μL of 0.5M EDTA was added, placed on ice for 3 min, then 2.5 μL of 1M NaOH and samples heated at 80℃ for 6 minutes, then held at 4℃. Then, 2.5 μL 1M HCl was added, and cleanup using the Oligo Clean & Concentrator kit was performed, eluting in 8 μL nuclease-free water. The cDNA was resolved in a 15% TBE-Urea polyacrylamide gel, and gel slices were resuspended in 600 μL DNA extraction buffer composed of 10 mM Tris-HCl pH 8.0, 300 mM NaCl, and 1 mM EDTA, then precipitated by freeze-thaw extraction as described above, resuspending in 8.5 μL nuclease-free water. For cDNA ligation, 1 μL of DMSO + 0.6 μL 2Puni linker (Table S3, 80 μM) were added to 7.5 μL cDNA and heated at 80℃ for 2 min, then held at 4℃. Then, an 11.5 μL master mix containing 2 μL 10X T4 RNA Ligase Buffer, 0.2 μL 100 mM ATP, 9 μL 50% PEG-8000, and 0.3 μL T4 RNA Ligase was added to each sample. After mixing, 1 μL of T4 RNA Ligase High Concentration (NEB, M0437M) was added to each reaction, then incubated overnight at 24℃, shaking at 1,600 rpm with 1 min on, 5 min off cycles, and then purified using the Oligo Clean & Concentrator kit, eluting in 35 μL nuclease-free water. Dual ligation products were amplified by PCR in a 50 µL reaction using Q5 High-Fidelity 2X Master Mix (NEB, M0492S) with Illumina indexing primers, then cleaned up with SPRI bead purification according to the manufacturer’s instructions (Bulldog Bio, CNGS050) and eluted in 13 μL nuclease-free water. Final libraries were gel purified on 4% E-Gel EX Agarose Gels (ThermoFisher, G401004), then quantified using a Qubit High Sensitivity dsDNA assay (Thermo Fisher Scientific, Q33231) with a Qubit fluorometer, as well as on a TapeStation (Agilent) with the High Sensitivity DNA assay ScreenTape (Agilent, 5067-5584). Final libraries were sequenced using 50 bp paired-end reads on an Illumina NovaSeq SP sequencer.

### Detection of ribosomal RNA modifications

To detect rRNA 2’-*O*-methyl and pseudouridine levels, we applied RiboMethSeq^66^ and bisulfite-induced deletion sequencing (BID-seq).^65,67^ Specifically, library preparation was performed as previously described^45^ with the following modifications: RNA was initially fragmented at 91℃ for 3 min. Following 3’ end ligation of RNA adaptor to RNA, each sample was split equally into a ‘+BID’ sample for pseudouridine detection and a ‘-BID’ sample for 2’-*O*-methyl detection and as a control for pseudouridine detection. -BID samples were frozen at -80℃ or placed on ice while +BID samples underwent bisulfite treatment and desulfonation as described previously.^65,67^ Reverse transcription into cDNA with SuperScript IV was then immediately performed on both -BID and cleaned, bisulfite-treated +BID samples with the following per-sample reaction described previously^67^: 4 µL 5X SuperScript IV RT buffer (ThermoFisher, 18090050), 2 µL 10 mM dNTP mix (NEB, N0447L), 1 µL 100 mM DTT (ThermoFisher, D1532), 0.5 µL RNase inhibitor (NEB, M0314L), 1 µL SuperScript IV (200 U) (ThermoFisher, 18090050), and 0.5 µL water. Reverse transcription was performed at 50℃ for 1 hr to allow for optimal pseudouridine detection. RNA-sequencing was performed on a NovaSeq 6000 (Illumina) using paired-end reads (150x150).

### Quantitative image analysis

#### Quantification of nucleolar core/rim enrichment of ribosome signal

Measurement of core and rim enrichment was as described previously^45^. Briefly, CellProfiler (v4.2.6)^113^ was used to automate image analysis. Nuclei labeled with SiR-DNA (Cytoskeleton Inc, CY-SC007) were segmented with Cellpose (v2.3.2)^114^ with an expected diameter of 150 pixels and the nuclei model. The distribution of mTagBFP2-NPM1 (nucleolar marker) was masked by the nuclei objects and used to segment nucleoli with pixel diameters between 15 and 200 using a global minimum cross-entropy threshold. Each nucleolus was then divided into 20 equal sized bins with the MeasureObjectIntensityDistribution module. The core was defined as the inner 3 bins and the rim as the outermost 3 bins and enrichment is the ratio of measured intensity divided by the expected intensity with a uniform distribution.

#### Quantification of cytoplasmic ribosome signal

Cytoplasmic ribosome signal quantification was performed using CellProfiler. The Hoechst (ThermoFisher, 62249) nuclear channel was first smoothed with a Gaussian filter (σ = 5) to smooth high-frequency features. Nuclei were segmented using a global minimum cross-entropy threshold, retaining objects with diameters between 100 and 250 pixels. Cell boundaries were then defined using theIntentifySecondaryObjects module, using the nuclei as input propagation objects, and global minimum cross-entropy thresholding. The nuclear region was then removed from the cell segmentation to produce a cytoplasmic-only mask which was used to measure cytoplasmic ribosome intensity in the cytoplasm of each cell.

#### Ribosome turnover analysis

Here, we quantify the decay of labeled ribosome concentration due to both degradation and dilution (from cell growth). The total amount of labeled ribosomes per cell (as measured by HaloTag-TMR intensity) at time 𝑡 is given by 𝑟_𝑙𝑎𝑏𝑒𝑙_ (𝑡) = 𝑟_0_ 𝑒^−*t*^, where the total decay rate (1/τ), 𝑘_𝑑𝑒𝑐𝑎𝑦_ = 𝑘_𝑑𝑒𝑔_ + µ is the sum of the degradation rate 𝑘_𝑑𝑒𝑔_ and the dilution rate µ. The dilution rate is estimated from the cell confluency, which is proportional to cell number 𝑛(𝑡) = 𝑛_0_ 𝑒*^ut^*. These two measurements gives the degradation rate as 𝑘_𝑑𝑒𝑔_ = 𝑘_𝑑𝑒𝑐𝑎𝑦_ − µ.

To obtain the degradation rate 𝑘_𝑑𝑒𝑔_ and the dilution rate µ, each time series (ribosome TMR label intensity and cell confluency) was modeled as a single exponential,

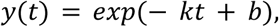

where 𝑘 is the rate constant and 𝑏 is the log-amplitude. Parameters were estimated by weighted nonlinear least squares, minimizing

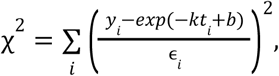

with ɛ*_i_* set to the SEM of each point. The parameters were initialized using a weighted linear fit to 𝑙𝑜𝑔(𝑦), with weights set by σ_𝑙𝑜𝑔𝑦_ ≈ ɛ*_i_* /*𝑦_i_*, and then optimized using Gauss–Newton iterations with an analytic Jacobian. The parameter uncertainties were estimated from a rescaled covariance matrix:

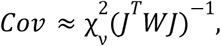

where 𝐽 is the Jacobian of parameters (𝑘, 𝑏) with respect to *X*^2^, 𝑊 is the diagonal weight matrix with entries 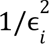, and 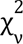 = *X*^2^ /(𝑑. 𝑜. 𝑓. − 2) is the reduced chi-squared. Standard errors were taken as the square roots of the covariance diagonal.

Degradation rates were computed as

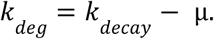

Uncertainties were propagated assuming independent errors,

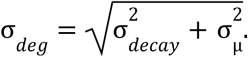

Reported errors correspond to one standard error of the fitted rate constants.

### Analysis of sequencing data

#### Ribo-seq data processing

FASTQ files were processed using the RiboFlow^115^ and RiboToolkit^116^ pipelines. Within the RiboFlow workflow, sequencing adapters were trimmed and low-quality reads were filtered using Cutadapt.^117^ Processed reads were aligned to the reference transcriptome using Bowtie2, and reads mapping to noncoding RNAs, etc. were filtered. Deduplication was performed using umi_tools. Ribosome footprint quality was assessed by metagene analysis from RiboFlow, confirming robust 3-nt periodicity. RiboToolkit was used to calculate translation efficiencies, transcript-level codon composition metrics and membrane protein terms. Translationally regulated transcripts were defined as those with a log₂ fold change ≥ 1.2 between molecularly aged (T72) and steady-state (T0) ribosomes and an adjusted p value < 0.05.

#### Ribosome pausing analysis

Identification of age-dependent ribosome pausing sites were performed as described previously.^32^ Coverage from riboflow (v0.0.1) was dumped for each sample using ribopy.^115^ Coding sequence (CDS) regions were isolated from each coverage map, and read counts were summed for each codon across biological replicates. Genes with an average coverage below 0.5 or fewer than 64 total reads were excluded from further analysis. Differential ribosome pausing was evaluated for each codon using a two-sided Fisher’s exact test comparing read counts between ‘young’ and ‘old’ samples. For the PROTAC experiment (Related to Figure 5), Control was treated as the ‘young’ condition and PROTAC treatment as the ‘old’ condition. Resulting p-values were adjusted for multiple testing using the Benjamini-Hochberg false discovery rate procedure. Significant sites were further filtered to remove positions where the ‘old’ count was below the gene-wide average, and where the ‘old’ samples exhibited fewer counts than ‘middle’ samples, when a ‘middle’ sample was available. Finally, codon sequences corresponding to each retained site were retrieved by indexing into the reference transcriptome. The resulting set represents codons exhibiting significantly increased ribosome pausing with molecular age. Amino acid enrichment at ribosomal E-, P-, and A-site positions was inferred by back-calculating site occupancy from mapped pause sites. Sequence logos were then generated using Seq2Logo v2.0.^118^

#### Ribosome occupancy analysis

Ribosome occupancy at each position was calculated from the riboflow coverage using a 20-codon window before and after the query codon. Occupancy was defined as the read count at each codon divided by the mean read count across the window. Query positions were derived either from age-dependent pausing sites or from polybasic regions. Polybasic K/R repeats were identified by translating each CDS transcript and scanning for stretches of three or more consecutive lysine (K) or arginine (R) residues. For each polybasic run, the start position, sequence, and corresponding occupancy profile were recorded.

#### Characterization of ribosome occupancy dynamics and identification of dropoff codons

The dynamics of ribosome occupancy during translation were estimated from ribosome footprint alignments generated by riboflow. For each gene of interest, pysam (v0.23.3) was used to compute per-base pileups across CDS regions, which were then aggregated into codon-level counts by summing reads across the three nucleotide positions. Counts were normalized to the maximum codon count per gene and sample. Ribosome occupancy profiles were subsequently smoothed using a triangular window spanning 11 codons (or 3 for data in Figure 5G) and normalized to the corresponding steady state sample counts at each codon position. Dropoff codons were identified by fitting relative ribosome occupancy profiles to a logistic function,

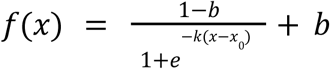

with the curve_fit function from scipy. The specific logistic function sets the maximum value to 1 with a variable minimum value, b which was clamped between 0 and 1; x_0_, the codon with a dropoff halfway between 1 and b was set to be > 0, and k, which controls the rate of dropoff, was left unbound. A gene was determined to have a dropoff if the logistic curve fit converged with a R2 score better than 0.4 and the parameters produced a reasonable codon for 90% and 50% dropoff from T0 (within the CDS). Codons upstream of the dropoff codon were used for further analysis. Sequence logos were generated using Seq2Logo v2.0.^118^

#### Analysis of ribosomal RNA modification levels

For 2’-*O*-methylation, 18S and 28S rRNA 2’-*O*-methylation levels from RNA-sequencing data was performed as previously described.^45^ Briefly, the fraction of RNA modified at known 2’-*O*-methylation sites was calculated by taking the weighted average of 5’ end read counts in ±2 nucleotide windows around each site (ScoreC). For pseudouridylation, Analysis of 18S and 28S rRNA pseudouridine levels from RNA-sequencing data was performed using a modified version of the published BID-pipe computational pipeline (https://github.com/SoftLivingMatter/pseudoU-BIDseq). Briefly, the ‘cutoff’ and ‘group-filter’ parameters were adjusted to allow calculation of pseudouridine modification levels for all rRNA nucleotides. As library preparation was reverse-stranded, the strandedness parameter was set to ‘false’. The ‘barcode’ string was set empty as no inline barcode was used. Paired-end reads (R1 and R2) were merged. Reads too short to be merged were treated as additional R1 reads, with R1 reads used as-is and R2 reads reverse-complemented prior to use. The fraction of RNA modified at known pseudouridine sites^65,67^ was used for further analysis and plotting.

#### Statistical determination of molecular age-dependent ribosomal RNA modifications

Monosome, disome, and unfractionated libraries each contained six groups defined by time point (T0 (steady state), T24, and T72 hours) and pulldown target (RPS17 or RPL10A), with two biological replicates per group. All analyses were performed in R (version 4.5.1). For each modification site, group differences were assessed using ANOVA from linear models (lm), with p-values adjusted for multiple testing using the Benjamini-Hochberg method (q < 0.05). Post hoc pairwise comparisons were conducted using Tukey’s HSD test, focusing on T0 vs. T72 hour contrasts within each pulldown; family-wise adjusted p-values < 0.05 were considered significant and were required to agree with the omnibus ANOVA result. Effect sizes were calculated as log₂(T72/T0) and are reported alongside Tukey-adjusted p-values in plots.

#### Statistical analysis

Where indicated, two-tailed Mann-Whitney U tests or Welch’s *t*-tests were applied as reflected in the figure legends, and were performed using GraphPad Prism 10. Normality was assessed where appropriate to guide test selection. Exact *p* values and sample sizes (n) are reported in the corresponding figure legends. Where indicated, gene ontology (GO) analysis was performed using the Gene Ontology Resource,^119^ with Bonferroni correction for multiple testing.

